# Dose-related Mutagenic and Clastogenic Effects of Benzo[b]fluoranthene in Mouse Somatic Tissues Detected by Duplex Sequencing and the Micronucleus Assay

**DOI:** 10.1101/2024.07.26.605228

**Authors:** D. M. Schuster, D. P. M. LeBlanc, G. Zhou, M. J. Meier, A. E. Dodge, P. A. White, A. S. Long, A. Williams, C. Hobbs, A. Diesing, S. L. Smith-Roe, J. J. Salk, F. Marchetti, C. L. Yauk

## Abstract

Polycyclic aromatic hydrocarbons (PAHs) are common environmental pollutants that originate from the incomplete combustion of organic materials. We investigated the clastogenicity and mutagenicity of benzo[*b*]fluoranthene (BbF), one of 16 priority PAHs, in MutaMouse males after a 28-day oral exposure. BbF causes robust dose-dependent increases in micronucleus frequency in peripheral blood, indicative of chromosome damage. Duplex Sequencing (DS), an error-corrected sequencing technology, reveals that BbF induces dose-dependent increases in mutation frequencies in bone marrow (BM) and liver. Mutagenicity is increased in intergenic relative to genic regions, suggesting a role for transcription-coupled repair of BbF-induced DNA damage. At higher doses, the maximum mutagenic response to BbF is higher in liver, which has a lower mitotic index but higher metabolic capacity than BM; however, mutagenic potency is comparable between the two tissues. BbF induces primarily C:G>A:T mutations, followed by C:G>T:A and C:G>G:C, indicating that BbF metabolites mainly target guanines and cytosines. The mutation spectrum of BbF correlates with cancer mutational signatures associated with tobacco exposure, supporting its contribution to the carcinogenicity of combustion-derived PAHs in humans. Overall, BbF’s mutagenic effects are similar to benzo[*a*]pyrene, a well-studied mutagenic PAH. Our work showcases the utility of DS for effective mutagenicity assessment of environmental pollutants.

**Synopsis:** We used Duplex Sequencing to study the mutagenicity of benzo[*b*]fluoranthene across the mouse genome. Dose-dependent changes in mutation frequency and spectrum quantify its role in PAH-induced carcinogenicity.

## Introduction

Identifying and characterizing the mutagenic properties of environmental chemicals is an essential component of human health risk assessment. Chemicals can cause somatic mutations or chromosomal aberrations that drive genetic diseases such as cancer (1,2). Presently, genetic toxicology studies are used for hazard identification; however, there is growing acceptance that mutations are endpoints of regulatory concern (3). Consequently, emphasis is now being placed on quantifying mutations with higher accuracy and precision, characterizing mechanisms of action, and conducting quantitative dose-response analyses of mutagenic endpoints.

Polycyclic aromatic hydrocarbons (PAHs) are ubiquitous environmental pollutants that pose health risks because of their reactivity with DNA (4). PAHs are formed through incomplete combustion of organic materials (5,6). Humans are exposed to PAHs through inhalation of diesel exhaust, asphalt fumes, combustion emissions such as tobacco smoke, ingestion of barbecued and smoked foods, or dermal contact. PAH exposure is linked to reproductive (7), cardiovascular (8), and other tissue toxicities (9,10). Most PAHs are classified by the International Agency for Research on Cancer (IARC) as probable (Group 2A) or possible (Group 2B) human carcinogens, with benzo[*a*]pyrene (BaP) being classified as carcinogenic to humans (Group 1) (11,12). Sixteen PAHs have been prioritized by the US Environmental Protection Agency due to their prevalence in the environment and known mutagenic and carcinogenic properties (13).

Benzo[*b*]fluoranthene (BbF) is a priority PAH that is associated with genotoxicity, carcinogenicity, adverse hormonal changes, kidney damage, and reproductive diseases (14,15,16,17). BbF is metabolized into reactive epoxides by cytochrome P450 (CYP) enzymes (18,19,20). Previous studies have shown that BbF is mutagenic in multiple tissues (10,21); however, these studies analysed mutagenesis in transgenic reporter genes. The effects of BbF on endogenous DNA and its mutational mechanisms are not well characterized.

Modern next-generation sequencing (NGS) technologies paired with error-correcting processes provide opportunities to comprehensively characterize mutagenesis (22). Duplex Sequencing (DS) is a targeted error-corrected NGS technology that allows the detection of rare mutations (23,24,25,26). It reduces the technical error rate of conventional NGS by uniquely tagging both strands of the original DNA fragments during library building, enabling the development of consensus sequences for each strand to eliminate PCR and sequencing errors (22,27,28). Importantly, DS quantifies mutation frequencies while in parallel providing detailed information on types of mutations, location, and clonal expansion. Information on target nucleotides can be used to link chemically induced trinucleotide mutation patterns to human cancer signatures to inform potential disease relevance (23,28,29).

Here, we used DS to study the mutagenicity of BbF in mouse liver, a metabolically active tissue with a moderate mitotic index, and bone marrow (BM), a tissue with low metabolic activity that is highly proliferative. We paired DS with the micronucleus assay to provide complementary information on the clastogenic effects of BbF in the same mice. The objectives were to: 1. quantify the dose-dependent effects of BbF on chromosome damage and mutation frequency; 2. determine the spontaneous and BbF-induced mutation spectra; 3. characterize variations in the mutagenic response between somatic tissues and across 20 genomic regions; 4. extract the BbF trinucleotide signature and identify similarities to the catalogue of somatic mutations in cancers (COSMIC); and 5. compare tissue- and endpoint-specific BbF mutagenic potency.

## Methods

### 1. Animal husbandry and chemical exposure

MutaMouse animals were maintained and exposed at Health Canada. The study was approved by the Health Canada Ottawa Animal Care Committee and followed the Canadian Council on Animal Care guidelines. Eight to ten weeks old males were exposed to doses of 0, 6.25, 12.5, 25, 50, or 100 mg/kg body weight/day (hereafter mg/kg/day) BbF (CAS 205-99-2; Sigma-Aldrich, Oakville, ON Canada) dissolved in olive oil for 28 consecutive days via oral gavage. Each dose group contained eight animals and the doses were selected based on a previous study that investigated the genetic toxicity of nine PAHs, including BbF (10).

### 2. Micronucleus (MN) assay

Two days following completion of chemical administration, we collected blood (N = 8 per group) from the saphenous vein using a Microvette® capillary tube coated with EDTA. For the MN assay, 200 µL of peripheral blood was processed according to the MicroFlow^PLUS-M^ Kit (Litron Laboratories, Rochester, NY, USA). Briefly, blood was diluted in heparin and fixed in ultra-cold methanol (Sigma-Aldrich, St. Louis, MO, USA). Samples were stored at -80°C for a minimum of three days before performing a buffer wash and stabilizing the fixed samples in the Long-Term Storage Solution of the MicroFlow^PLUS-M^ Kit. The peripheral blood was then shipped to Integrated Laboratory Systems (ILS), an Inotiv Company (North Carolina, USA), on cold packs. Eight samples per dose group were labeled with anti-mouse CD71-FITC and anti-rodent CD61-PE antibodies and run on a BD FACSCalibur™ dual-laser bench top low cytometer. The assay detects micronucleated reticulocytes (MN-RET) and red blood cells (MN-RBC) as per Organisation for Economic Co-operation and Development (OECD) Test Guideline 474 (30). At least 20,000 ± 2,000 RET and approximately 1,000,000 RBC were enumerated per sample. The frequencies of MN-RET and MN-RBC were expressed per 1000 cells, and the %RET was used as an indicator of BM toxicity. We performed a pairwise comparison to the vehicle control using the Bonferroni multiple comparison test, with P < 0.05 being significant.

### 3. Tissue collection and DNA extraction

BM and liver tissue were harvested from euthanized animals three days after the end of exposure as per the OECD test guideline 488 (31). The liver was separated into its right posterior, right anterior, median, caudate, and left lobe. We collected BM by centrifugation of the femurs after removing the proximal and distal ends, and the cell pellet was then resuspended in phosphate-buffered saline (Corning™ cellgro™, Manassas, VA, USA) (32). Liver and BM samples were flash frozen and stored at −80 °C until analysis.

DNA was extracted from the right anterior liver lobe and BM of four randomly selected animals per dose group. The extractions were performed using the Qiagen DNeasy blood and tissue kit protocol (Cat. # 69504, Qiagen, Hilden, Germany) with modifications. Specifically, tissue disruption by bead beating the samples twice at 4000 rpm for 30 seconds was added, and proteinase K digestion was conducted with a decreased temperature of 37°C and an increased incubation duration of 180 minutes. The extracted DNA was analysed using the Qubit™ 1x dsDNA Broad Range Assay Kit (Thermo Fischer, Waltham, MA, USA) to determine DNA concentration and the Agilent TapeStation Genomic ScreenTape assay (Agilent Technologies, Santa Clara, CA, USA) to evaluate its integrity. All samples had DNA Integrity Numbers > 7.0 and concentrations > 25 ng/µL as required for DS library preparation (23).

### 4. DS library preparation and sequencing

Library preparations for targeted DS were performed using the TwinStrand Duplex Sequencing® Mutagenesis Panel (Mouse-50) v1.0 (MMP) according to the DS manual provided by TwinStrand Biosciences (33). Briefly, 500 ng of DNA was enzymatically fragmented to a final size of approximately 300-400 base pairs (bp). Fragments were then end-polished, poly A-tailed, and ligated to DS adapters with unique molecular identifiers (UMIs) (TwinStrand Biosciences, Seattle, WA, USA). Quantification of the library concentrations was performed using Qubit™ 1x dsDNA High Sensitivity Assay Kits (Thermo Fisher, Waltham, MA, USA). The Agilent TapeStation High Sensitivity D1000 Screen Tape assay (Agilent Technologies, Santa Clara, CA, USA) was used to determine the average fragment sizes. The mouse mutagenesis panel (MMP) is designed to capture 20 endogenous loci (2.4 kb each) that are distributed across all mouse autosomes, with chromosome one containing two loci. The loci were selected by TwinStrand Biosciences to broadly represent the nucleotide composition, GC content, heterochromatin state, and transcriptional properties of the mouse genome (24). Repetitive regions and pseudogenes were excluded to ensure efficient hybrid capture. Cancer driver genes or genes under strong negative selective pressure were also omitted. Of the 20 loci, nine are located within genic and eleven within intergenic regions according to the Mouse GenCode gene database version M25 (34).

Sequencing was performed at Psomagen (Rockville, MD 20850, USA) on a NovaSeq 6000 S4 flow cell, targeting a minimum of 600 million raw reads passing filters per sample with at least 10,000-fold target locus coverage. All samples surpassed minimum sequencing threshold; all libraries produced over 700 million informative duplex-bases, mean on-target sequencing higher than 10,000x, and sufficient peak tag families. Sequencing data are found in the Sequence Read Archive (SRA) under SRA accession number PRJNA1131313. These FASTQ files were processed using the TwinStrand DS Mutagenesis App version 3.4.2 (TwinStrand Biosciences, Seattle, WA, USA) (23). This application extracts the 12-nucleotide barcode sequence and removes the 5 bp adapter from the read sequence. After checking for non-nucleotide characters, the barcode sequence is parsed to create a duplex tag header. After paired-end read alignment and sorting, a single strand consensus sequence (SSCS) is produced *via* ConsensusMaker.py. Afterwards, SSCS are sorted and analyzed *via* DuplexMaker.py to obtain the double strand consensus sequence (DSCS). This data processing uses UMIs to identify and eliminate technical errors and false mutation discoveries. The error-correction reduces the error rate to less than one error per 10^−7^ to 10^−8^ nucleotides, enabling the detection of rare mutagenic events (35).

### 5. DS Data Interpretation and Statistical Analysis

We further processed the DS data to calculate the tissue-specific mutation frequency (MF). First, we identified and counted each mutation by aligning the sequencing data to the reference genome of *Mus musculus* (mm10, GRCm38.p6). Since cells carrying mutations can clonally expand, tissues may contain multiple identical mutations that arose from a single mutagenic event. We first calculated mutation frequencies under the conservative assumption that identical mutations arose from the same mutagenic event. Under this assumption, all identical mutations within a sample were counted as one single mutation to calculate the minimum mutation frequency (MF_min_). We also counted all mutations to obtain the maximum mutation frequency (MF_max_) (25). The variant allele fraction was set to 0.01 to exclude mosaicism and germline mutations.

MF_min_ and MF_max_ estimations by dose were calculated using R software based on generalized linear models with an over-dispersed binomial error distribution. The doBy R package was used to perform pairwise comparisons and the estimates were back-transformed (36). We calculated the back-transformed standard error using the delta method, and a Holm-Sidak correction was applied to adjust the p-values for multiple comparisons. Binomial error distribution of generalized linear mixed models (GLMM) was used to calculate the locus-specific MFs and pairwise comparisons were made using the doBy R package as described above (36).

We obtained the single base spectrum by direct comparison of unique substitution mutations post data processing to the mm10 reference genome. Mutation counts were summed across dose group using the SigProfiler Matrix Generator (37). The dose and tissue specific 96-trinucleotide signature were determined by considering the adjacent nucleotides located upstream and downstream of single base substitutions. These trinucleotide signatures were then reconstructed using single base substitution signatures (SBS) of the Catalogue of Somatic Mutations in Cancer (COSMIC) database with the SigProfiler Assignment tool (29). The accuracy metrics for this reconstruction include the calculation of cosine similarity between the chemically induced trinucleotide pattern and the reconstructed spectrum (38). In general, the threshold for cosine similarities is set to 0.8 with cosine similarities greater than 0.9 being considered a reliable reconstruction (39).

### 6. Benchmark Dose Modeling and potency evaluation

Benchmark dose (BMD) modeling was performed using the PROAST v70.1 web tool on the MN-RET frequencies, MN-RBC frequencies, and the MF_min_ values for liver and BM. The value for the Akaike Information Criterion (AIC), a measure of the goodness of fit for each model, was used to identify best fit models. A difference of 2 units was used to indicate that one model had a better fit than another following EFSA recommendations (40). BMD calculations were fit to the 3- and 5-parameter Hill, Exponential, Inverse Exponential, and Log-Normal models. The most probable BbF dose-response curve for the various endpoints was determined by averaging 200 bootstrap runs. The AIC was calculated throughout all bootstrap runs and unfit models were excluded from the model averaging. We used a 50% increase compared to the vehicle control as the benchmark response (BMR) (41,42,43). The respective BMD lower and upper confidence limit values (i.e., BMDL and BMDU) were derived from 90% confidence intervals.

## Results

We evaluated the effect of repeat-dose, 28-day oral exposure to BbF on clastogenicity and mutagenicity in MutaMouse males. BbF-induced chromosome damage was measured using the flow-cytometric peripheral blood MN assay. MF and spectral changes caused by BbF were quantified using DS. We note that no significant changes in body and organ weights were detected in BbF exposed animals (data not shown).

To quantify the extent to which BbF exposure causes chromosome damage, we measured MN-RET and MN-RBC frequencies in peripheral blood collected two days following the 28-day exposure. BbF caused a dose-dependent increase in the frequencies of MN-RET and MN-RBC (Figure 1, Supplementary Table 1), with significant increases (P < 0.05) in all dose groups relative to olive oil controls. MN-RET frequencies per 1000 cells increased from 2.34 ± 0.1 in the control group to 8.11 ± 0.4 in the high dose group (100 mg/kg/day), and MN-RBC frequencies increased from 1.63 ± 0.1 in the control group to 4.84 ± 0.2 in the high dose group. We also observed a significant increase from 1.60 ± 0.1 to 2.17 ± 0.1 in %RET (Supplementary Table 1), which is indicative of BbF-induced BM toxicity (44). These results provide evidence of BM toxicity and a robust increase in chromosome damage caused by oral exposure to BbF.

**Figure 1.**
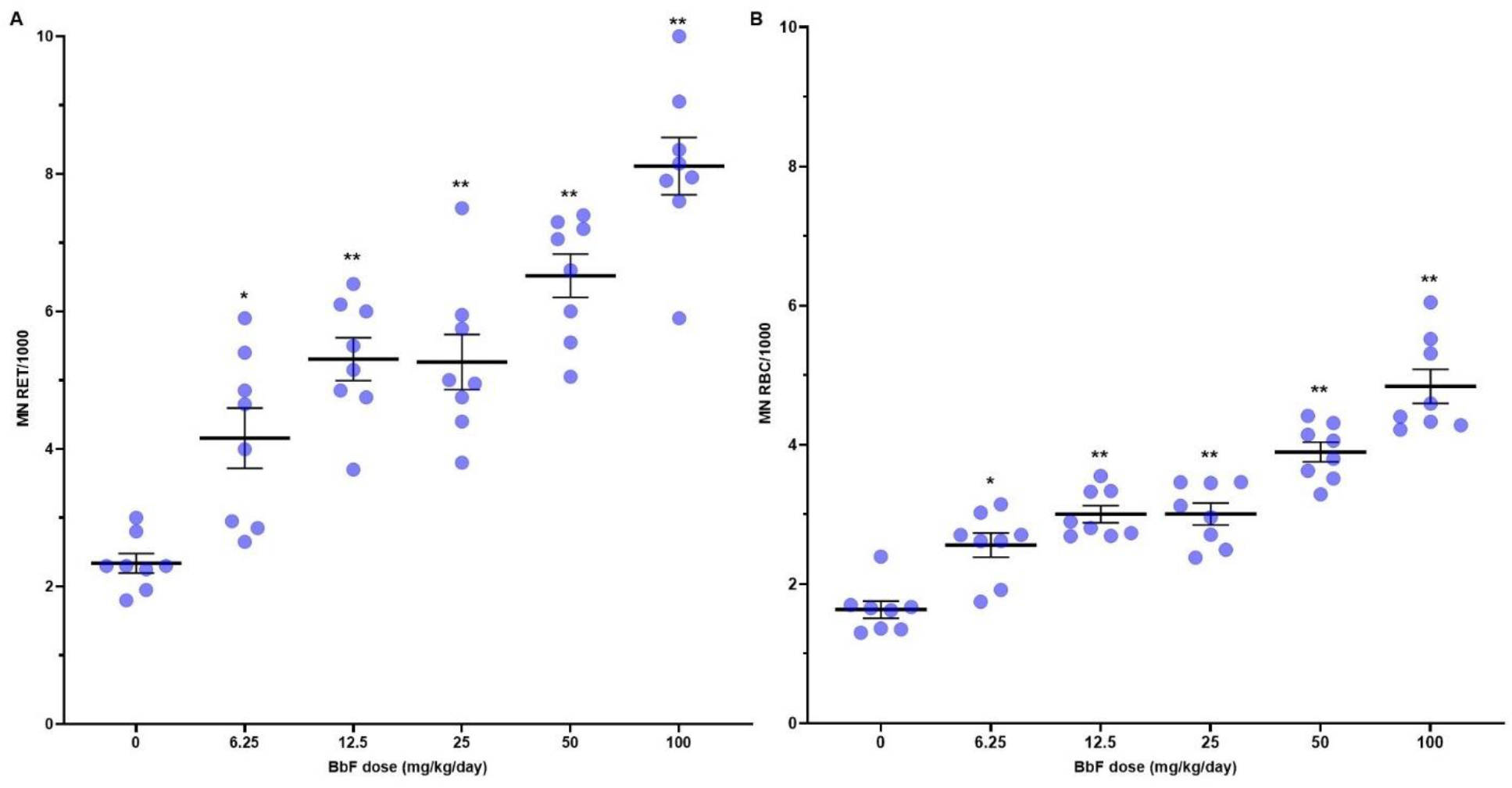
Frequencies of spontaneous and chemically induced micronuclei in peripheral blood of MutaMouse males after BbF exposure for 28 days. Mean micronucleus (MN) frequencies (black bar) with indication of standard errors of the mean (SEM). MN frequencies per 1000 cells are shown for both reticulocytes (MN RET) (A) and red blood cells (MN-RBC) (B). Dots represent individual animal data. ^*^ P < 0.05 and ^**^ P < 0.01 relative to vehicle control.

Next, we applied DS to a panel of 20 endogenous loci to investigate the mutagenic impacts of BbF exposure. Our sequencing recovered between 2.21 × 10^+8^ to 7.06 × 10^+8^ raw reads passing filter per sample in BM, and 2.10 × 10^+8^ to 6.81 × 10^+8^ in liver (Supplementary Table 2). On average, there were approximately 1 billion informative duplex bases per sample in both tissues and DS reads were distributed evenly across the 20 target regions. Single nucleotide variants (SNVs), insertions and deletions (indels), and multi-nucleotide variants (MNVs) were quantified. An average of 36, 37, 64, 66, 98, and 131 unique mutations were identified in BM and 40, 48, 79, 88, 196, and 327 in liver for 0, 6.25, 12.5, 25, 50, and 100 mg/kg/day BbF dose groups, respectively.

Our analysis reveals that oral exposure to BbF causes robust and statistically significant dose-dependent increases in MF_min_ in BM and liver (Figure 2). Spontaneous MF_min_ (× 10^−8^ ± SEM) was 4.13 ± 0.45 in BM and 5.33 ± 1.01 in liver, respectively. MF_min_ increased to 14.78 ± 0.76 in BM and 36.27 ± 2.49 in liver at the high dose (Figure 2, Supplementary Table 3). The fold change relative to controls in the top dose was greater in liver (6.8-fold increase) than in BM (3.6-fold increase). Significant increases in MF_min_ were found for all but the lowest dose in BM and the two lowest doses in liver.

**Figure 2.**
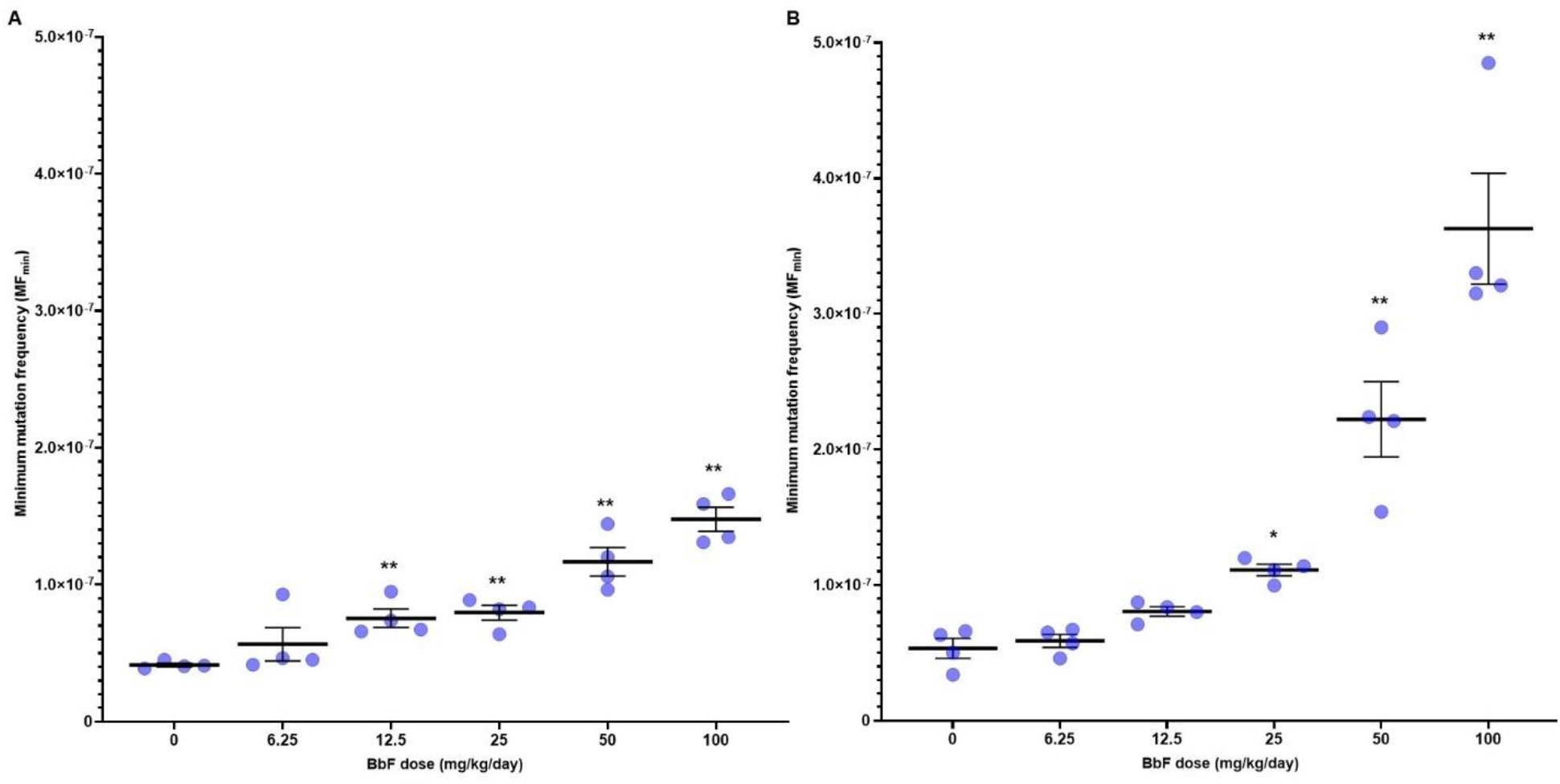
Mean minimum mutation frequency (MF_min_ ± SEM) per bp measured by Duplex Sequencing in mouse bone marrow (A) and liver (B) after exposure to increasing doses of BbF (N = 4 per dose group). Asterisks indicate ^*^ P < 0.05 and ^**^ P < 0.01 relative to vehicle control. Dots represent individual animal MF_min_.

To explore the extent of clonal expansion of cells carrying mutations, we examined the DS data in each individual animal to quantify all mutagenic events (MF_max_). MF_max_ was consistently greater than MF_min_ in each animal in both tissues (Supplementary Table 4). In BM, there were two animals with notable amounts of clonal expansion, i.e., animals 6 and 10 in the 6.25 and 12.5 mg/kg/day dose groups, respectively (Figure 3a). Clonal expansion in these animals was driven by multiple clonally expanded mutations rather than being caused by one or two mutational events. In contrast, there were no major instances of clonal expansion events in liver (Figure 3b). Moreover, animals 6 and 10 did not have the same clonal expansion patterns in liver as were observed in BM. Finally, there was higher inter-animal variability in MF_max_ compared to MF_min_ (Figure 3a and 3b). These results indicate that clonal expansion events in our study were tissue-specific and relatively low in frequency.

**Figure 3.**
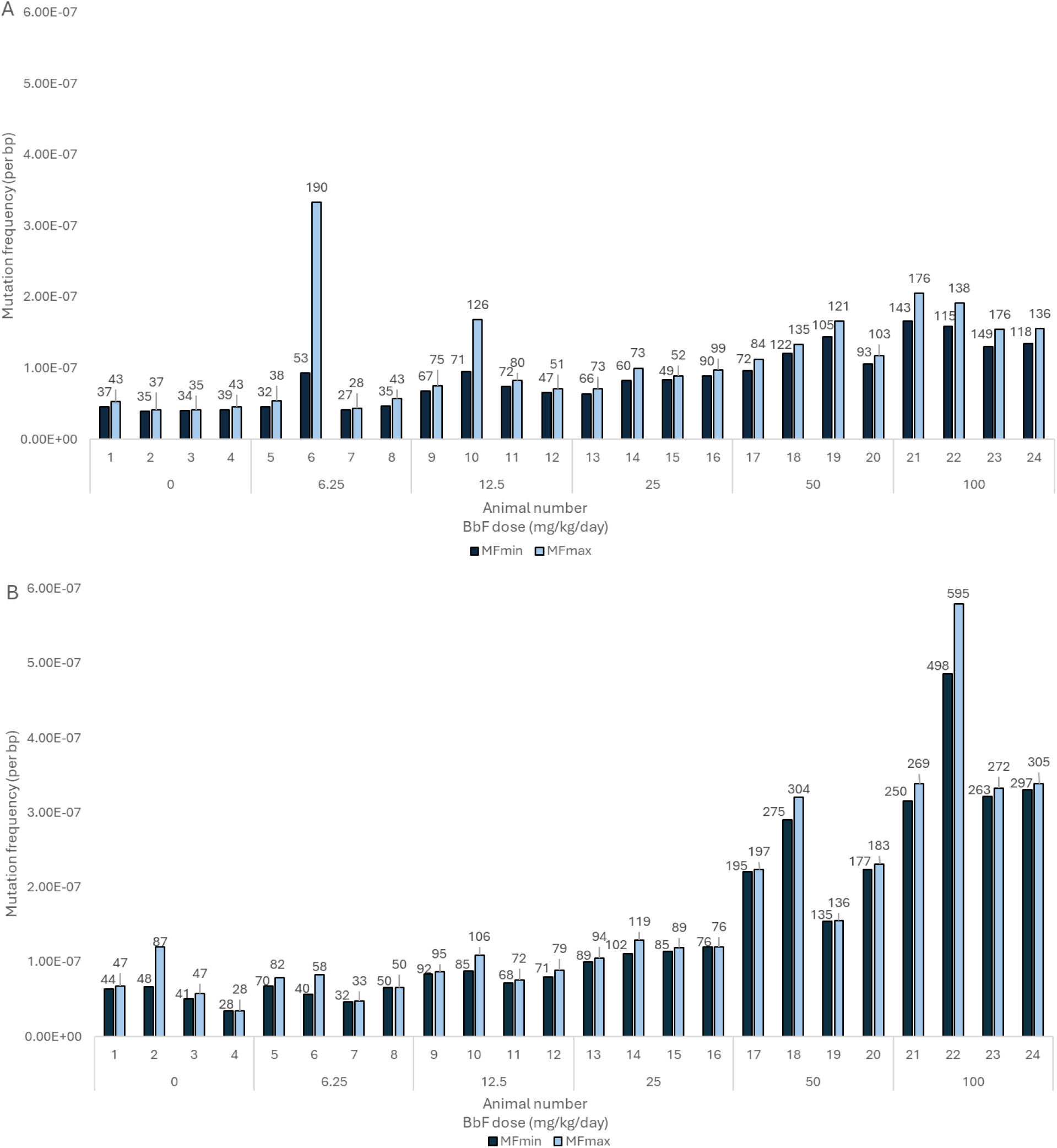
Mean mutation frequency per bp measured by Duplex Sequencing in bone marrow (A) and liver (B) for individual mice within each BbF dose group. The mutation frequency of unique mutations is indicated as MF_min_ (black). The mutation frequency including clonally expanded mutations is shown as MF_max_ (light blue). Numbers above the bars represent the observed number of mutations.

Next, we investigated whether there are locus-specific differences in spontaneous and BbF-induced mutagenicity across the MMP loci. In BM, there was a statistically significant dose-dependent increase in MF_min_ in 9 out of 20 loci relative to vehicle controls (i.e., with MF_min_ per locus at the top dose being significantly higher than vehicle control) (Figure 4a). The top five most BbF-responsive loci at the highest dose are on chromosomes 14, 11, 8, 17, and 16, with MF_min_ of 37.50 ± 3.58, 35.18 ± 6.74, 34.73 ± 10.20, 34.70 ± 3.78, and 30.38 ± 6.63 × 10^−8^. The lowest Mf_min_ of 3.80 ± 0.72 × 10^−8^ occurred in the locus on chromosome 3 (Figure 4a, Supplementary table 5). There is a 9.9-fold difference in MF_min_ at the top dose between the most and least responsive loci in BM.

**Figure 4.**
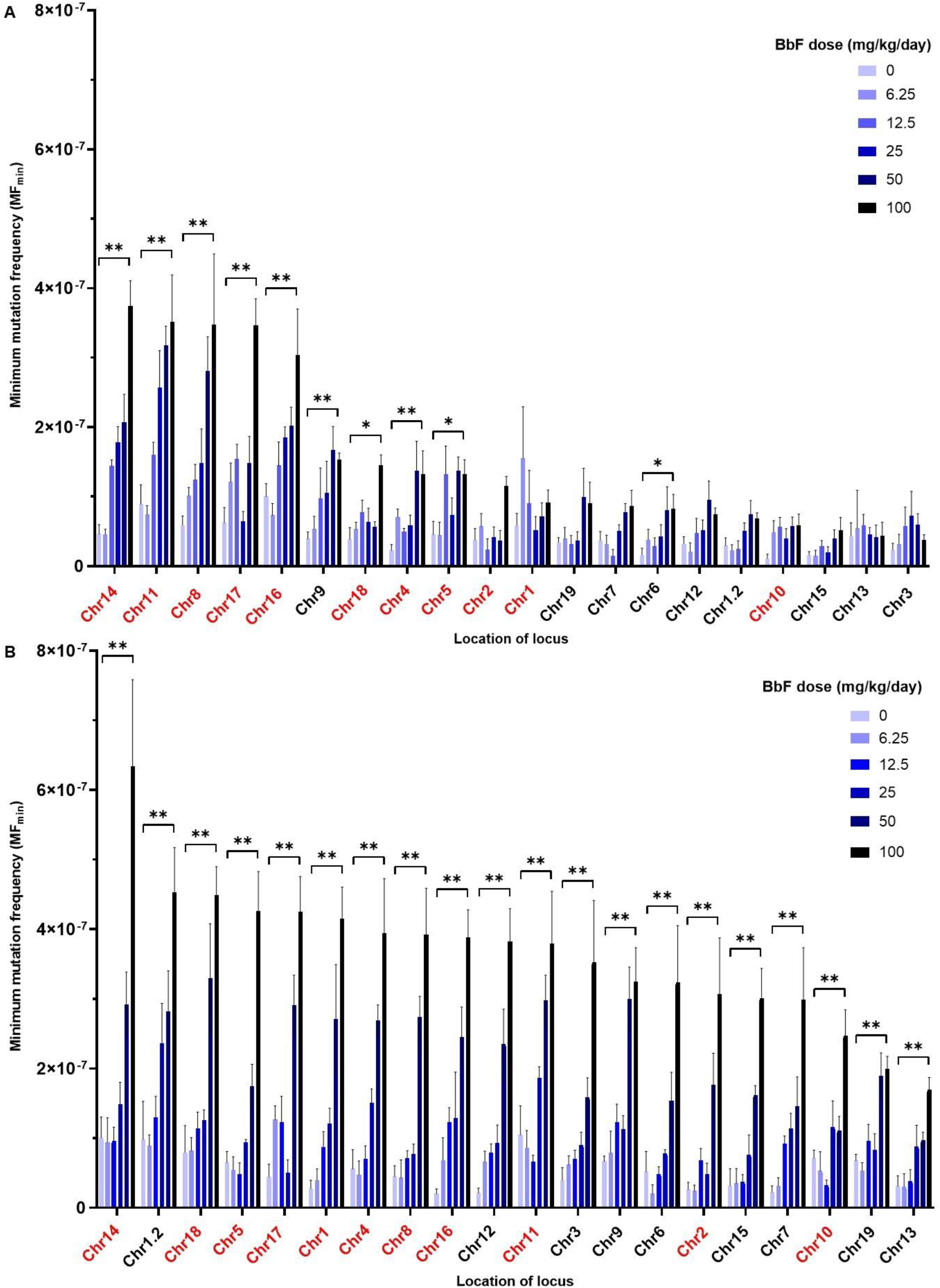
Mean minimum mutation frequency (MF_min_) ± SEM per locus for BbF-exposed mouse bone marrow (A) and liver (B) alongside vehicle controls. Loci are sorted from the highest top dose response to the lowest (left to right). Intergenic and genic loci are show in red and black, respectively. Statistical testing was done for the top dose with respect to the vehicle control using a generalized linear mixed model; ^*^ P < 0.05 and ^**^ P < 0.01; N = 4 per dose group.

In liver, all loci exhibited significant dose-dependent increases in MF_min_ following BbF exposure (Figure 4b). At the top dose, the top 5 most responsive loci are located on chromosomes 14, 1.2, 18, 5, and 17, with MF_min_ of 63.35 ± 12.50, 45.28 ± 6.47, 44.88 ± 4.12, 42.63 ± 5.68, and 42.50 ± 5.09 × 10^−8^, respectively. There was a 3.7-fold difference in MF_min_ between the most and least responsive loci in liver. The locus on chromosome 13 had the lowest MF_min_ (17.00 ± 1.72 × 10^−8^) (Supplementary table 6). As in BM, the locus located on chromosome 14 has the highest MF.

Overall, the majority of most BbF-responsive loci in both tissues are intergenic. The average MF_min_ was 21.8 and 40.5 × 10^−8^ in intergenic loci, and 6.7 and 31.2 × 10^−8^ in genic loci, in BM and liver respectively. The difference between intergenic and genic loci was significant between the two and the four highest doses in BM and liver, respectively. Overall, our results indicate that the observed susceptibility to BbF-induced mutations is locus-specific.

DS enables the detailed characterization of chemically induced mutation subtypes to provide insight into underlying mutagenic mechanisms. To understand the mutation spectrum induced by BbF, we analysed SNVs, categorized into single base substitution types according to the pyrimidine reference code, indels and MNVs. C:G>T:A was the most common mutation subtype in controls, with a MF_min_ of 2.34 ± 0.24 × 10^−8^ in BM and 4.75 ± 0.57 × 10^−8^ in liver. BbF induced dose-dependent increases in C:G>A:T and C:G>T:A mutations compared to the vehicle control (P < 0.01 at the top dose for BM and liver), with highly similar spectra observed in the two tissues (Figure 5; Supplementary Table 7). C:G>A:T substitutions had the highest MF_min_ at the top dose reaching 12.78 ± 1.41 × 10^−8^ in BM (6.1-fold above control) and 35.48 ± 5.62 × 10^−8^ in liver (29.3-fold above control). C:G>T:A MF_min_ increased to 7.84 ± 0.81 (3.3-fold) and 16.53 ± 1.37 × 10^−8^ (3.5-fold) in BM and liver, respectively. Overall, the BbF mutation spectrum is consistent between tissues and is mainly driven by C:G>A:T and C:G>T:A mutations.

**Figure 5.**
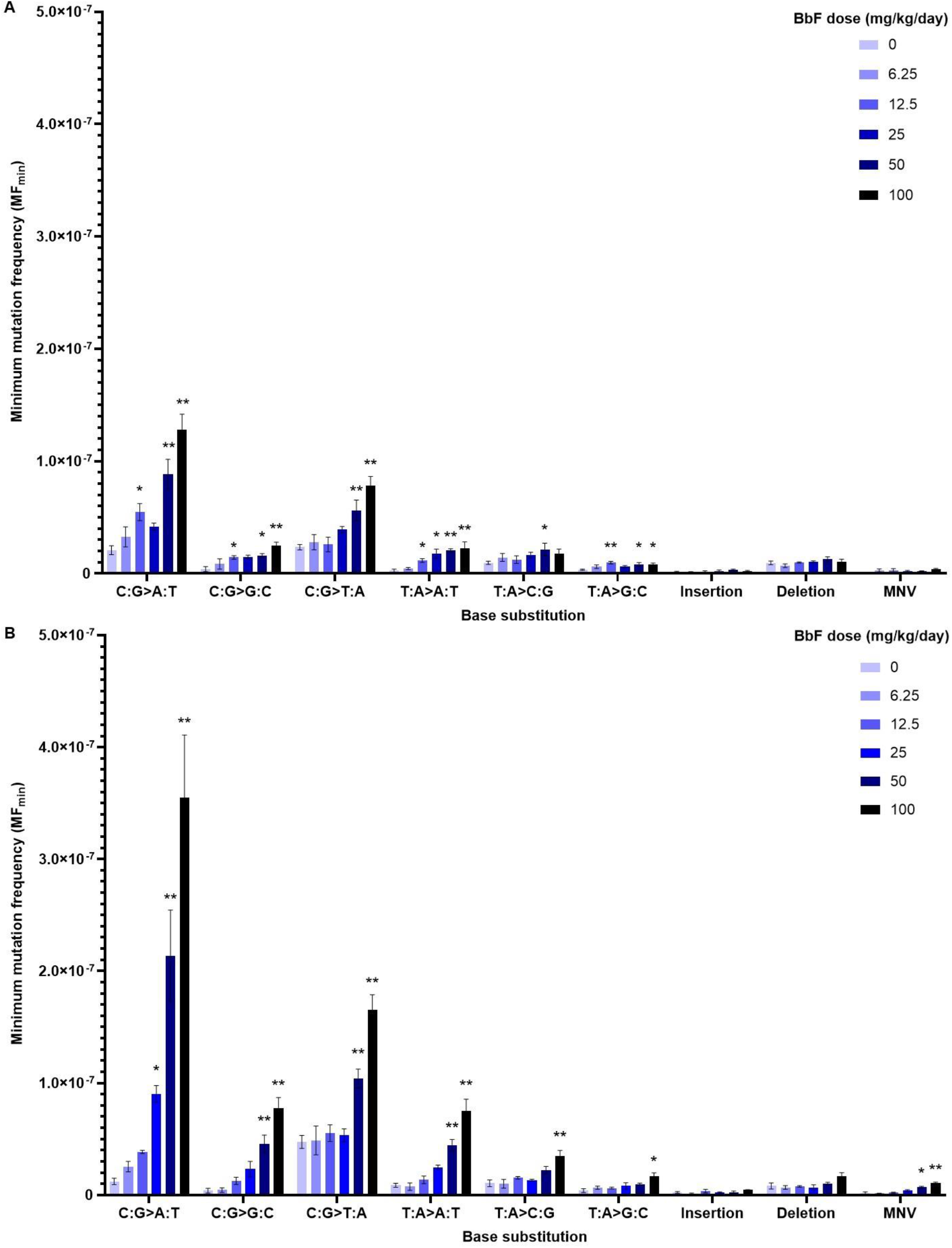
Mutation spectrum for vehicle controls and BbF dose groups observed in mouse bone marrow (A) and liver (B). Mutation subtypes, insertions, deletions and multi-nucleotide variants (MNV) are presented as mean MF_min_ ± SEM. Mutation subtypes are based on the six single nucleotide variants using a pyrimidine reference. Statistical testing was done using a generalized linear mixed model; ^*^ P < 0.05 and ^**^ P < 0.01; N = 4 per dose group.

MF_min_ for insertions in the vehicle controls was 0.12 ± 0.08 × 10^−8^ in BM and 0.24 ± 0.07 ×10^−8^ in liver. Insertion MF_min_ at the top BbF dose increased to 0.21 ± 0.07 and 0.45 ± 0.03 × 10^−8^ for BM and liver, respectively. Deletions occurred in the vehicle control with frequencies of 0.95 ± 0.17 × 10^−8^ in BM and 0.81 ± 0.27 × 10^−8^ in liver. Deletion MF_min_ at the top BbF dose was 1.03 ± 0.26 and 1.65 ± 0.33 × 10^−8^ for BM and liver, respectively. The detected increases in indels were not statistically significant in both tissues. The frequencies of MNVs increased from 0.06 ± 0.03 to 0.41 ± 0.04 × 10^−8^ in BM and from 0.14 ± 0.14 to 1.06 ± 0.09 × 10^−8^ in liver reaching statistical significance (p = 0.004; Supplementary Table 7).

We characterized the trinucleotide mutation spectrum to determine the influence of adjacent nucleotides on BbF-induced mutagenicity. The control samples were enriched in C:G>T:A mutations with the highest frequencies observed in the ACA, CCA, and TCA trinucleotide context. BbF caused a shift in the trinucleotide spectrum driven predominantly by increases in C:G>A:T mutations at CCT, CCA, ACA, and TCT trinucleotides (Supplementary Figure 1). We used the SigProfiler toolset to reconstruct the trinucleotide signature of vehicle controls and BbF-treated tissues. COSMIC signature SBS5 was enriched in control BM (25%) and liver (36%) (Supplementary Figure 2); SBS5 is associated with ageing and is found in most cancers and in normal cells. Exposure to BbF shifted the trinucleotide mutation pattern in BM to be associated with SBS29 (32%), which originates from tobacco chewing, and SBS5 (19%). The reconstructed signature had a cosine similarity of 0.94 with the original signature (Figure 6a). The enriched signatures in liver at the top dose were SBS4 (75%), which is associated with tobacco smoking, and SBS5 (14%). The liver reconstructed signature had a cosine similarity of 0.96 with the original signature (Figure 6b). SBS4 showed a dose-dependent increase in liver mutation profiles; no other SBS showed dose-dependent increases. Our data demonstrate dose-dependent changes in the trinucleotide mutation pattern with enrichment of cancer signatures SBS4, and SBS29.

**Figure 6.**
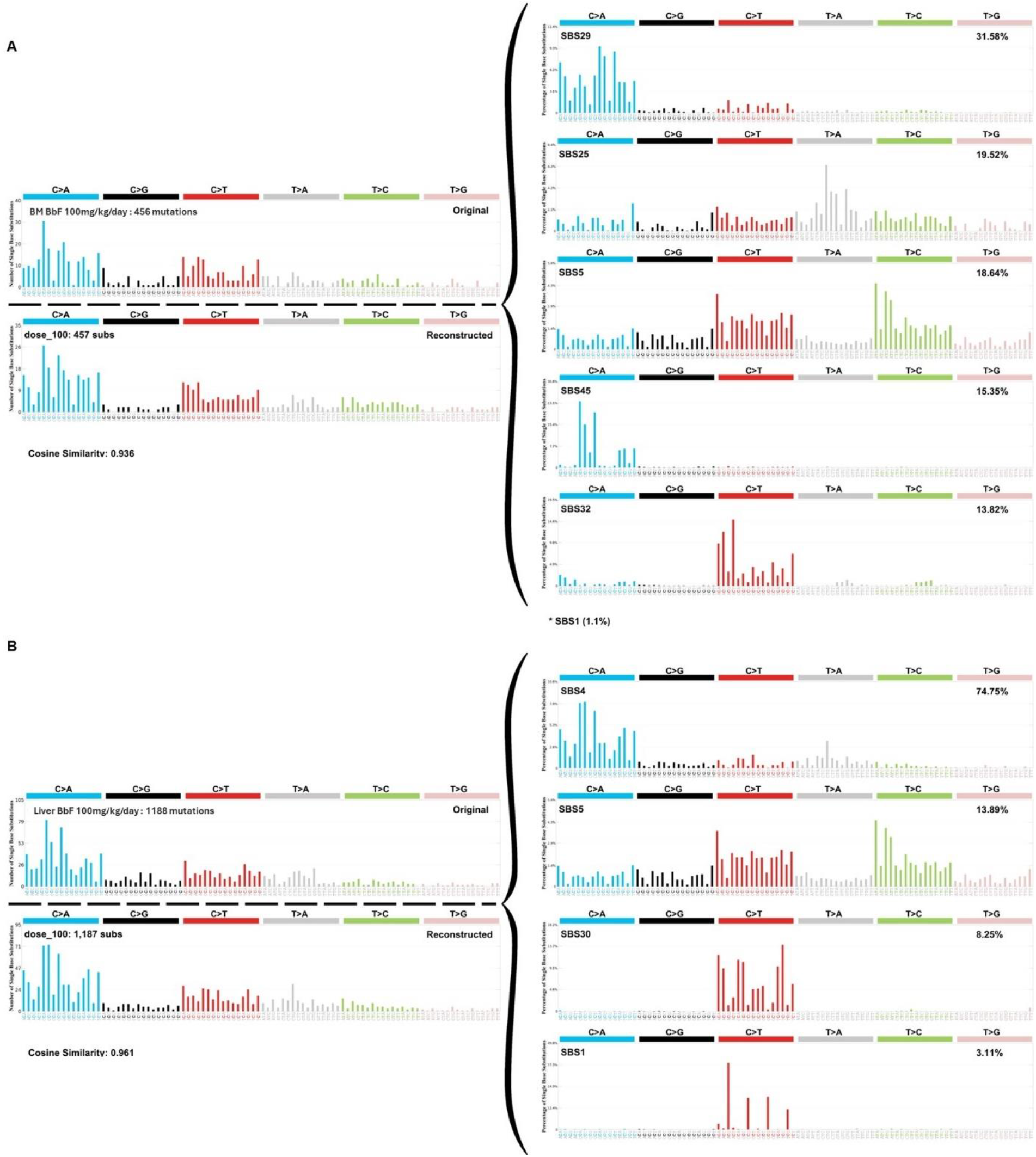
Mutation signature analyses for top BbF dose (100 mg/kg/day) in bone marrow (A) and liver (B). The original trinucleotide mutation profile is shown on the top left. SigProfilerAssignment used the single base substitution (SBS) signatures of the Catalogue of Somatic Mutations in Cancer (COSMIC) database to reconstruct the signatures. The SBS signatures and their relative contributions are shown on the right. The reconstructed mutation pattern, number of substitutions used for the assignment, and cosine similarity between reconstructed and observed trinucleotide mutation profile are shown on left below the original profile. The total number of mutations in the original signature are indicated on the left.

We employed BMD modeling to compare the potency of BbF in inducing clastogenicity and mutagenicity across endpoints and tissues. The BMDs are presented as confidence intervals (BMDL – BMDU plotted in order of increasing BMDs) for comparison (Figure 7). The BMD confidence interval for MF_min_ in BM was overlapping with all other confidence intervals, indicating comparable BbF potencies between the endpoints. However, the BMD confidence intervals for MN-RET and MN-RBC were overlapping but were both lower than the confidence interval for MF_min_ in liver (Figure 7, Supplementary Table 8). Thus, a 50% increase in clastogenicity in RET and RBC occurs at lower doses than a 50% increase in liver MF_min_. Confidence interval ranking from least sensitive to most sensitive endpoint/tissue follows the pattern: MF_min_ in liver (10.1 to 18.4 mg/kg/day), followed by MF_min_ in BM (3.6 to 12.8 mg/kg/day), then MN frequency in RBCs (2.2 to 9.3 mg/kg/day), and finally MN frequency in RET (0.4 to 4.1 mg/kg/day). Our results show that the BMD confidence intervals were generally within a comparable range, indicating that there are no major tissue- or endpoint-specific differences in mutagenic potencies of BbF.

**Figure 7:**
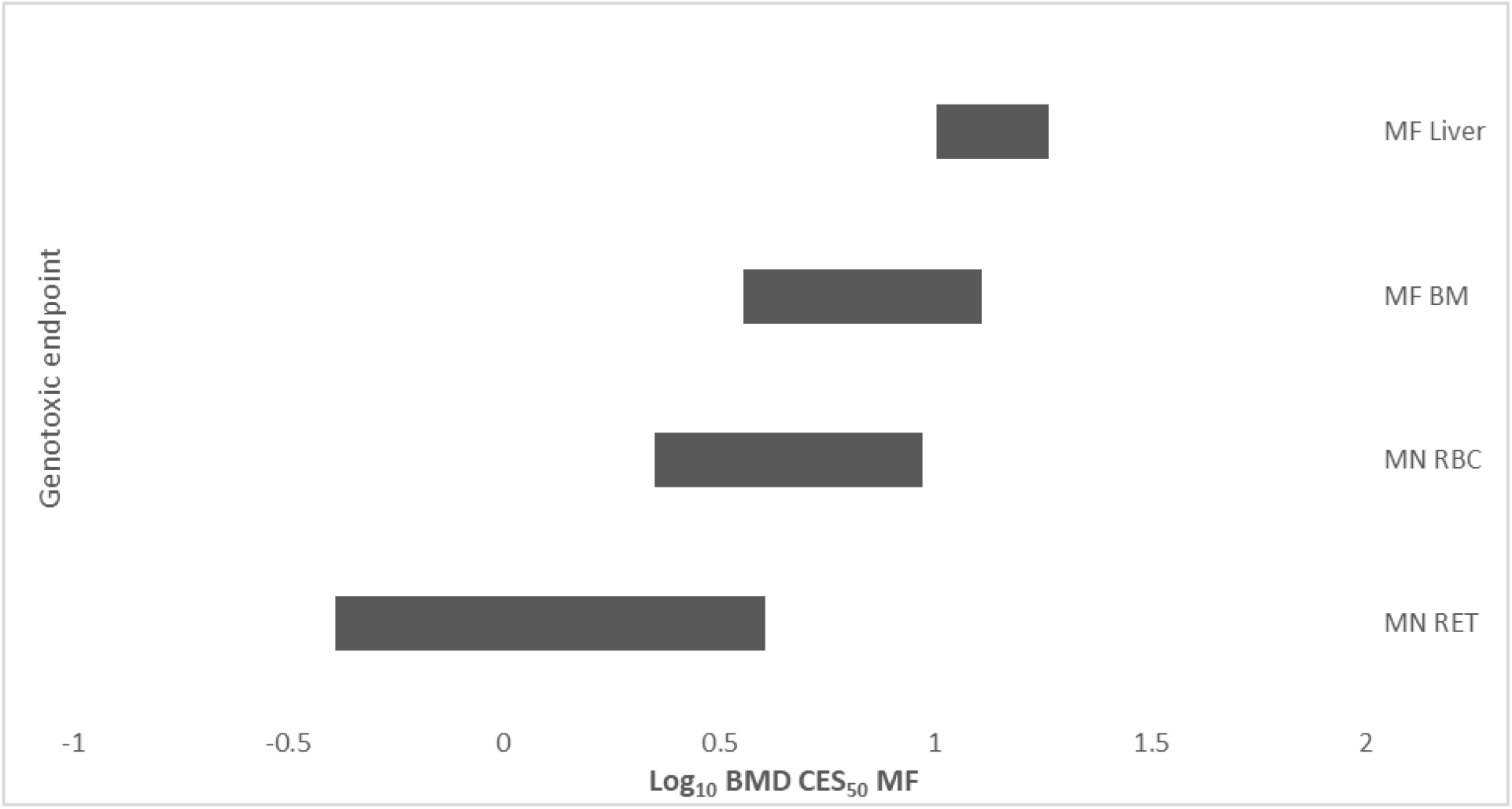
Log_10_ benchmark dose (BMD) for a 50% increase in micronucleus (MN) or mutation frequency (MF_min_). 90% confidence intervals based on BMD model averaging analyses using PROAST are shown. Confidence intervals are organized from lowest (top) to highest (bottom) potency. Labels indicate micronuclei in reticulocytes (MN-RET) and red blood cells (MN-RBC), and mutation frequency in liver (MF liver) and bone marrow (MF BM).

## Discussion

We investigated the effects of sub-chronic oral exposure of MutaMouse males to BbF on mutagenicity and clastogenicity. As expected, our results show that BbF causes dose-dependent increases in MF and MN. The magnitude of the mutagenic response was larger in liver than in BM at the highest dose tested; however, dose-response modeling to estimate the BMD50 for MF_min_ reveals overlapping confidence intervals (i.e., similar potencies) for the two tissues. Greater increases in MF due to BbF exposure were observed in intergenic than genic loci, consistent with transcription-coupled repair of bulky adducts. BbF induces mostly C:G>A:T mutations with a spectrum similar to other PAHs such as BaP (24). Mutational signature analysis following this short-term exposure shows a correlation between BbF-induced mutations in both liver and BM and signatures observed in human cancers associated with tobacco exposure. The data support that BbF contributes to the carcinogenicity of PAH mixtures in humans and underscores the human health significance of exposure to this PAH. Our work also showcases the high value of DS in deeply characterizing the mutagenic effects of exposure to environmental genotoxicants.

We observed robust increases in chromosome damage in blood cells and mutagenic responses in liver and BM. Our MN analysis extends the findings of Long et al. (10) to demonstrate significant clastogenic effects at lower BbF doses. Long et al. also showed a higher mutagenic response in liver *lacZ* reporter gene mutations than in BM, which is consistent with our findings; however, our analyses quantify the mutagenicity of BbF to endogenous loci rather than a single exogenous bacterial reporter gene, increasing the biological relevance of the findings to human health. Indeed, our data demonstrate large differences in BbF response across the 20 loci in the MMP, supporting the need to look beyond individual genes. The liver is more metabolically active, proliferates more slowly, and has a lower DNA replication rate than BM. Although more reactive metabolites are created in the liver than in BM, DNA damage has more time to be repaired prior to replication and has less opportunity to be converted into mutations during the exposure period (44). At the same time, despite the lower metabolic activity, the high proliferation rate makes the hematopoietic system susceptible to MN formation. Our results indicate systemic distribution and strong clastogenic and mutagenic effects after oral exposure to BbF that reflect the metabolic capacity and proliferation rate of the two tissues.

To further compare the response between the two tissues and endpoints, we used BMD modeling to identify the doses at which BbF exposure elicits a 50% increase in MN or MF. The BMD confidence intervals for MN-RETs and MN-RBCs, and mutations in BM, overlapped. The only notable difference was that MF in liver was slightly lower than that for MN-RETs. Thus, we observe no major differences in potencies for these tissues/endpoints. Interestingly, the mutagenicity potency of BbF in BM is comparable to the mutagenic potency of BaP in BM assessed using the MMP (24). The confidence intervals span 3.6 to 12.8 mg/kg/day and 4.5 to 7.5 mg/kg/day for BbF and BaP, respectively. Overall, our results indicate that BbF is a potent genotoxicant in both hematopoietic and liver tissues and that its mutagenic potency in BM is comparable to that of BaP, which is the frame of reference when evaluating PAHs.

Similarly to previous studies (23,24,25), our analyses revealed major differences in response to BbF across the 20 loci of the mouse mutagenesis panel, with MF_min_ being higher in intergenic than genic loci. Two of these studies applied the same MMP used in our study. The top responding loci are nearly identical across these studies, which analyzed chemicals with different mutagenic mechanisms. LeBlanc et al. (24) studied the bone marrow of MutaMouse males exposed for 28 days to BaP. BbF and BaP, especially their hydrodiol and quinone metabolites, cause bulky DNA adducts that target guanines or cytosines and are mainly repaired by nucleotide excision repair (NER); thus, consistencies across these exposures may be expected (18,46). In contrast, Dodge et al. (25) measured mutations following exposure to procarbazine, which causes DNA alkylation that is restored by base excision repair (BER) (47,48). All three chemicals also cause an increase in reactive oxygen species that can oxidize DNA, which is then repaired by BER (49,50,). While NER and BER can both be coupled to active transcription, PAH adducts are mainly repaired by TC-NER (51,52). Our study is consistent with the hypothesis that higher induced MFs in intergenic regions might be driven by the lack of TCR activity in these loci (23,25). In contrast, the same genic versus intergenic trend has not been observed in DS experiments with Sprague-Dawley rats. These studies administered N-nitrosodiethylamine and N-ethyl-N-nitrosourea, which are common alkylating agents (26,53). More research across different model systems and chemicals with different mechanisms of action will be required to effectively determine the underlying drivers of these intriguing locus-specific differences in mutagenic responses.

One of the powerful attributes of DS is its ability to directly quantify the extent of clonal expansion of any mutation. Thus, for each locus we compared MF_min_ and MF_max_. MF_min_ is highly informative for characterizing the potency and mutational signature of a mutagen. MF_max_ has physiological importance for the organism since mutated cells that clonally expand statistically increase the likelihood of multiple clonal events within a cluster of cells carrying the same mutation that may render a cell population more prone to tumour formation (54,55). However, we note that the MMP loci were chosen in part because they are not thought to be under positive or negative selective pressure; thus, clonal expansion should be a random event. Herein, we observed no major differences between MF_min_ and MF_max_ in most of the samples. A key factor for clonal expansion of mutations is time (56). The 28-day exposure duration in adult mice may be too short for notable clonal expansion to occur within the observed loci. With increasing exposure duration or sampling time, we may expect to see an increase in clonal expansion especially in rapidly proliferating tissues like BM. Overall, the low levels of clonal expansion of BbF-induced mutations does not significantly affect MF in the tissues studied, but differentiation between unique and clonal mutations is valuable for understanding the mutational burden in each tissue.

The application of DS also enabled an in-depth analysis of the mutation spectrum induced by BbF to provide insight into its mechanism of action. In line with expectations, and similarly to BaP, BbF causes C:G>A:T and C:G>T:A mutations. Although the specific nucleophilic sites of BbF adduct formation are not known (10), previous studies have identified the amino group of guanines as the most likely target for PAH-DNA adduct formation through reacting with diol-epoxides or hydroquinones (57,16). The metabolism of BbF is expected to create these highly reactive intermediates that can engage in oxidation-reduction, nucleophilic addition, and electrophilic aromatic substitution reactions (58). The liver is the main xenobiotic metabolizing tissue and has high CYP levels, with CYP1A1 and CYP1B1 isoforms being key enzymes for PAH metabolism (59,60). However, due to the high instability of BbF metabolites, it is unlikely that they are transported from the liver to the BM unless they are stabilized within the blood serum (61). Thus, we postulate that BbF requires metabolism in the BM through local CYP enzymes such as the highly abundant CYP1B1 (62). Overall, our results support that both tissues metabolize BbF. The consistencies in mutation spectrum in both tissues and similarities to BaP strongly suggest that BbF metabolites predominantly form cytosine or guanine adducts and that BbF is systemically distributed.

The trinucleotide mutational spectrum provides deeper insights into underlying biological processes associated with mutagenesis; it can be used to explore similarities with human cancer signatures. We observed tissue-specific differences in the trinucleotide patterns in exposed samples. Both tissues showed dose-dependent increases in C:G>A:T mutations, but the upstream adjacent nucleotides differed. In BM, mutations were found predominantly in a G<u>C</u>N>G<u>A</u>N context; whereas, in liver they were predominantly in the C<u>C</u>N>C<u>A</u>N context. Signature reconstruction indicated that SBS29, associated with oral tobacco exposure, was the strongest contributing signature in BM; while in liver, SBS4, which is associated with tobacco smoking and is also associated with the DS mutational spectrum of BaP (24), was the highest contributing signature. SBS29 is enriched in mutations in a G<u>C</u>A>G<u>A</u>A, G<u>C</u>C>G<u>A</u>C, or G<u>C</u>T>G<u>A</u>T context, whereas SBS4 consists of C<u>C</u>A>CAA, C<u>C</u>C>C<u>A</u>C, and C<u>C</u>T>C<u>A</u>T variants. The respective aetiologies of these signatures align with common routes of BbF exposure for humans since chewing tobacco (63) and tobacco smoke (64) contain high levels of BbF. Both signatures are mainly driven through mutations in intergenic regions. The detection of these cancer-related signatures after 28 days of BbF exposure emphasizes the relevance of BbF to human PAH-induced carcinogenesis.

BbF administration over the time course of 28 days for this project aligns with OECD test guideline 488 (31), which is considered the ‘gold standard’ for regulatory mutagenicity assessment. Our study confirms that this short-term exposure is also sufficient to produce a robust DS dose-response in endogenous mouse loci in both of the tissues studied. This study and others support that DS mutagenicity assessment can be integrated with MN frequency assessment in 28-day studies to comprehensively characterize chromosome damage and mutagenic responses in vivo (23,24,30). Our results also suggest that DS could be integrated with other conventional toxicology studies such as the OECD test guideline 407 (65). Such integration would enable an evaluation of mutagenicity and clastogenicity within the standard 28-day repeated dose studies in rodents to significantly reduce animal use in toxicity testing, aligning with the 3Rs principles (66).

In conclusion, our study provides detailed insight into the mutagenic and clastogenic characteristics of BbF and its underlying mechanisms of action in the mammalian genome. In addition to dose-dependent increases in MN and MF in tissues with different metabolic activity and mitotic indices, we showed that C:G>A:T mutations are the drivers for BbF’s mutational signature. We observed enrichment of mutational signatures found in human PAH-associated cancers in both tissues after a 28-day BbF exposure. This finding underscores the significant role of BbF in environmental carcinogenesis. Finally, our findings highlight the high value of DS in mutagenicity assessment of chemical exposures and its potential utility for regulatory testing.

## Acknowledgements

CLY, FM, and PAW hold an Innovation in Regulatory Science Award from the Burroughs Wellcome Fund (#1021737). CLY is grateful for funding from the Natural Sciences and Engineering Research Council of Canada (RGPIN-2021-02806) with infrastructure support through the Canadian Foundation for Innovation John Evans Leaders Fund (#233109), the Ontario Research Fund (#40569), and the University of Ottawa. CLY acknowledges that this project was conducted, in part, thanks to funding provided through the Canada Research Chairs Program (CRC-2020-00060). Work at Health Canada was supported by the Chemicals Management Plan to FM. FM and MM acknowledge funding from Health Canada’s Genomics Research and Development Initiative. SSR was funded by the intramural research program of the National Institutes of Environmental Health Sciences, National Institutes of Health (ES103378-01) and genetic toxicity testing conducted by Integrated Laboratory Systems, an Inotiv company, under NIEHS contract 75N96020C00001. We thank Dr. Scott S. Auerbach at the Division of Translational Toxicology, NIEHS, and Marc Beal from Health Canada for their helpful review of the manuscript.

## Disclaimer

Health Canada does not endorse or recommend the products or services of any commercial entity, including TwinStrand Biosciences, Inc.

