## Supplementary Data for "Dose-related Mutagenic and Clastogenic Effects of Benzo[b]fluoranthene in Mouse Somatic Tissues Detected by Duplex Sequencing and the Micronucleus Assay"

Supplementary Table 1: Peripheral blood micronucleus assay conducted 2 days after 28 days of oral exposure to BbF.

| BbF Dose (mg/kg/day) | n | MN-RET[1]/1000 ± SEM | Adj. P value[2] | MN-RBC[3]/1000 ± SEM | Adj. P value[2] | %RET ± SEM | Adj. P value[2] |
|---|---|---|---|---|---|---|---|
| 0 | 8 | 2.34 ± 0.1 | - | 1.63 ± 0.1 | - | 1.60 ± 0.1 | |
| 6.25 | 8 | 4.16 ± 0.4 | 0.0036 | 2.56 ± 0.2 | 0.0014 | 1.81 ± 0.1 | 0.5763 |
| 12.5 | 8 | 5.31 ± 0.3 | <0.0001 | 3.00 ± 0.1 | <0.0001 | 1.87 ± 0.1 | 0.2245 |
| 25 | 8 | 5.26 ± 0.4 | <0.0001 | 3.00 ± 0.2 | <0.0001 | 2.02 ± 0.1 | 0.0146 |
| 50 | 8 | 6.52 ± 0.3 | <0.0001 | 3.90 ± 0.1 | <0.0001 | 2.24 ± 0.1 | <0.0001 |
| 100 | 8 | 8.11 ± 0.4 | <0.0001 | 4.84 ± 0.2 | <0.0001 | 2.17 ± 0.1 | 0.0005 |

[1] Micronucleated reticulocytes.
[2] Pairwise comparison to 0 mg/kg/day BbF using the Bonferroni multiple comparison test, significant if P < 0.05.
[3] Micronucleated red blood cells.

Supplementary Table 2: Technical performance metrics of the libraries built on DNA from BM and liver from animals after exposure to BbF.

| BbF Dose (mg/kg/day) | Animal Number | Raw Reads Passing Filter | Median Insert Size | Mean Duplex Depth | Peak Tag Family Size | Informative Duplex Bases | Raw Reads Passing Filter | Median Insert Size | Mean Duplex Depth | Peak Tag Family Size | Informative Duplex Bases |
|---|---|---|---|---|---|---|---|---|---|---|---|
| | | Bone Marrow | | | | | Liver | | | | |
| 0 | 1 | $3.12^{+08}$ | 299 | 15,550 | 14 | $1.17^{+09}$ | $2.46^{+08}$ | 252 | 16,764 | 13 | $1.04^{+09}$ |
| 0 | 2 | $3.01^{+08}$ | 269 | 17,271 | 11 | $1.34^{+09}$ | $6.81^{+08}$ | 267 | 17,433 | 40 | $1.07^{+09}$ |
| 0 | 3 | $3.25^{+08}$ | 285 | 15,948 | 14 | $1.22^{+09}$ | $2.53^{+08}$ | 276 | 19,600 | 11 | $1.18^{+09}$ |
| 0 | 4 | $3.73^{+08}$ | 249 | 18,109 | 13 | $1.47^{+09}$ | $2.73^{+08}$ | 259 | 19,867 | 12 | $1.23^{+09}$ |
| 6.25 | 5 | $3.42^{+08}$ | 292 | 13,438 | 19 | $1.00^{+09}$ | $5.19^{+08}$ | 268 | 25,014 | 20 | $1.53^{+09}$ |
| 6.25 | 6 | $7.06^{+08}$ | 425 | 10,718 | 63 | $7.06^{+08}$ | $2.10^{+08}$ | 248 | 16,886 | 9 | $1.05^{+09}$ |
| 6.25 | 7 | $3.02^{+08}$ | 311 | 12,297 | 19 | $9.08^{+08}$ | $2.78^{+08}$ | 276 | 16,524 | 18 | $9.60^{+08}$ |
| 6.25 | 8 | $2.40^{+08}$ | 278 | 14,593 | 9 | $1.11^{+09}$ | $2.43^{+08}$ | 261 | 18,443 | 10 | $1.13^{+09}$ |
| 12.5 | 9 | $3.20^{+08}$ | 271 | 19,131 | 10 | $1.47^{+09}$ | $4.99^{+08}$ | 261 | 26,398 | 19 | $1.63^{+09}$ |
| 12.5 | 10 | $5.42^{+08}$ | 308 | 14,064 | 29 | $1.07^{+09}$ | $3.70^{+08}$ | 248 | 23,489 | 13 | $1.49^{+09}$ |
| 12.5 | 11 | $3.48^{+08}$ | 276 | 18,558 | 12 | $1.43^{+09}$ | $4.72^{+08}$ | 261 | 23,011 | 20 | $1.44^{+09}$ |
| 12.5 | 12 | $2.30^{+08}$ | 284 | 13,785 | 10 | $1.04^{+09}$ | $3.55^{+08}$ | 272 | 21,224 | 16 | $1.28^{+09}$ |
| 25 | 13 | $3.56^{+08}$ | 276 | 19,780 | 11 | $1.52^{+09}$ | $3.14^{+08}$ | 268 | 21,397 | 13 | $1.30^{+09}$ |
| 25 | 14 | $2.21^{+08}$ | 279 | 14,154 | 9 | $1.06^{+09}$ | $4.34^{+08}$ | 286 | 22,006 | 20 | $1.33^{+09}$ |
| 25 | 15 | $3.34^{+08}$ | 319 | 11,071 | 22 | $8.31^{+08}$ | $3.34^{+08}$ | 267 | 17,970 | 18 | $1.10^{+09}$ |
| 25 | 16 | $4.57^{+08}$ | 220 | 19,177 | 15 | $1.62^{+09}$ | $2.65^{+08}$ | 202 | 15,613 | 9 | $1.10^{+09}$ |
| 50 | 17 | $2.40^{+08}$ | 273 | 14,280 | 11 | $1.10^{+09}$ | $2.64^{+08}$ | 268 | 21,146 | 10 | $1.28^{+09}$ |
| 50 | 18 | $3.71^{+08}$ | 251 | 19,331 | 12 | $1.55^{+09}$ | $3.12^{+08}$ | 255 | 22,785 | 12 | $1.39^{+09}$ |
| 50 | 19 | $4.03^{+08}$ | 294 | 13,726 | 21 | $1.04^{+09}$ | $3.60^{+08}$ | 254 | 21,148 | 16 | $1.31^{+09}$ |
| 50 | 20 | $3.02^{+08}$ | 268 | 16,705 | 12 | $1.27^{+09}$ | $4.06^{+08}$ | 275 | 19,002 | 21 | $1.17^{+09}$ |
| 100 | 21 | $3.38^{+08}$ | 230 | 16,505 | 9 | $1.39^{+09}$ | $3.40^{+08}$ | 286 | 19,003 | 17 | $1.15^{+09}$ |
| 100 | 22 | $2.38^{+08}$ | 266 | 13,861 | 11 | $1.06^{+09}$ | $3.80^{+08}$ | 253 | 24,706 | 14 | $1.54^{+09}$ |
| 100 | 23 | $4.56^{+08}$ | 282 | 21,661 | 14 | $1.67^{+09}$ | $2.76^{+08}$ | 236 | 19,811 | 11 | $1.26^{+09}$ |
| 100 | 24 | $2.85^{+08}$ | 271 | 16,844 | 10 | $1.30^{+09}$ | $4.81^{+08}$ | 249 | 21,772 | 21 | $1.39^{+09}$ |

Supplementary Table 3: Minimum mutation frequencies by duplex sequencing using the mouse mutagenesis panel in bone marrow and liver samples after exposure to increasing doses of BbF.

| BbF Dose (mg/kg/day) | n | $MF_{min}$[1] ± SEM (×$10^{-8}$) | Adj. P value[2] | $MF_{min}$ Fold-change[3] |
|---|---|---|---|---|
| | Bone Marrow | | | |
| 0 | 4 | 4.13 ± 0.45 | - | - |
| 6.25 | 4 | 5.65 ± 0.59 | 0.6271 | 1.4 |
| 12.5 | 4 | 7.54 ± 0.58 | 0.0052 | 1.8 |
| 25 | 4 | 7.95 ± 0.60 | 0.0025 | 1.9 |
| 50 | 4 | 11.66 ± 0.72 | <0.0001 | 2.8 |
| 100 | 4 | 14.78 ± 0.76 | <0.0001 | 3.6 |
| | Liver | | | |
| 0 | 4 | 5.33 ± 1.01 | - | - |
| 6.25 | 4 | 5.88 ± 1.05 | 0.9807 | 1.1 |
| 12.5 | 4 | 8.06 ± 1.10 | 0.3567 | 1.5 |
| 25 | 4 | 11.12 ± 1.43 | 0.0378 | 2.1 |
| 50 | 4 | 22.23 ± 1.94 | <0.0001 | 4.2 |
| 100 | 4 | 36.27 ± 2.49 | <0.0001 | 6.8 |

[1] Minimum mutation frequency considering unique mutations.
[2] Pairwise comparison to 0 mg/kg/day BbF by GLM, significant if $P < 0.05$.
[3] Fold-change of minimum mutation frequency versus 0 mg/kg/day BbF.

Supplementary Table 4: Minimum and maximum mutation frequencies in bone marrow and liver samples from individual animals after exposure to increasing doses of BbF.

| BbF Dose (mg/kg/day) | Animal | $MF_{min}$[1] ($\times 10^{-8}$) | $MF_{max}$[2] ($\times 10^{-8}$) | Fold-change[3] | $MF_{min}$[1] ($\times 10^{-8}$) | $MF_{max}$[2] ($\times 10^{-8}$) | Fold-change[3] |
|---|---|---|---|---|---|---|---|
| | | Bone marrow | | | Liver | | |
| 0 | 1 | 4.51 | 5.24 | 1.2 | 6.32 | 6.75 | 1.1 |
| 0 | 2 | 3.87 | 4.10 | 1.1 | 6.61 | 12.0 | 1.8 |
| 0 | 3 | 4.04 | 4.16 | 1.0 | 5.01 | 5.75 | 1.1 |
| 0 | 4 | 4.09 | 4.51 | 1.1 | 3.39 | 3.39 | 1.0 |
| 6.25 | 5 | 4.51 | 5.36 | 1.2 | 6.72 | 7.87 | 1.2 |
| 6.25 | 6 | 9.29 | 33.3 | 3.6 | 5.70 | 8.26 | 1.4 |
| 6.25 | 7 | 4.15 | 4.30 | 1.0 | 4.60 | 4.75 | 1.0 |
| 6.25 | 8 | 4.63 | 5.69 | 1.2 | 6.51 | 6.51 | 1.0 |
| 12.5 | 9 | 6.72 | 7.53 | 1.1 | 8.38 | 8.65 | 1.0 |
| 12.5 | 10 | 9.49 | 16.8 | 1.8 | 8.74 | 10.9 | 1.2 |
| 12.5 | 11 | 7.38 | 8.20 | 1.1 | 7.11 | 7.53 | 1.1 |
| 12.5 | 12 | 6.58 | 7.14 | 1.1 | 8.02 | 8.92 | 1.1 |
| 25 | 13 | 6.38 | 7.06 | 1.1 | 9.97 | 10.5 | 1.1 |
| 25 | 14 | 8.20 | 9.98 | 1.2 | 11.1 | 12.9 | 1.2 |
| 25 | 15 | 8.34 | 8.85 | 1.1 | 11.4 | 11.9 | 1.0 |
| 25 | 16 | 8.88 | 9.77 | 1.1 | 12.0 | 12.0 | 1.0 |
| 50 | 17 | 9.63 | 11.2 | 1.2 | 22.1 | 22.3 | 1.0 |
| 50 | 18 | 12.0 | 13.3 | 1.1 | 29.0 | 32.0 | 1.1 |
| 50 | 19 | 14.4 | 16.6 | 1.2 | 15.4 | 15.5 | 1.0 |
| 50 | 20 | 10.6 | 11.7 | 1.1 | 22.4 | 23.1 | 1.0 |
| 100 | 21 | 16.6 | 20.5 | 1.2 | 31.5 | 33.9 | 1.1 |
| 100 | 22 | 15.9 | 19.1 | 1.2 | 48.5 | 58.0 | 1.2 |
| 100 | 23 | 13.1 | 15.4 | 1.2 | 32.1 | 33.2 | 1.0 |
| 100 | 24 | 13.5 | 15.5 | 1.2 | 33.0 | 33.9 | 1.0 |

[1] Minimum mutation frequency considering unique mutations.
[2] Maximum mutation frequency considering all detected mutations.
[3] Fold-change between $MF_{min}$ and $MF_{max}$

Supplementary Table 5: Minimum frequencies per loci detected in bone marrow samples after exposure to increasing doses of BbF.

| | 0 mg/kg/day | 6.25 mg/kg/day | | | 12.5 mg/kg/day | | | 25 mg/kg/day | | | 50 mg/kg/day | | | 100 mg/kg/day | | |
|---|---|---|---|---|---|---|---|---|---|---|---|---|---|---|---|---|
| Location of locus | $MF_{min}$[1] ± SEM (x $10^{-8}$) | $MF_{min}$[1] ± SEM (x $10^{-8}$) | Fold-change[2] | Adj. P-Value[3] | $MF_{min}$[1] ± SEM (x $10^{-8}$) | Fold-change[2] | Adj. P-Value[3] | $MF_{min}$[1] ± SEM (x $10^{-8}$) | Fold-change[2] | Adj. P-Value[3] | $MF_{min}$[1] ± SEM (x $10^{-8}$) | Fold-change[2] | Adj. P-Value[3] | $MF_{min}$[1] ± SEM (x $10^{-8}$) | Fold-change[2] | Adj. P-Value[3] |
| Chr1 | 5.91 ± 1.69 | 15.51 ± 7.44 | 2.6 | 0.9986 | 9.07 ± 4.70 | 1.5 | 0.9994 | 5.14 ± 2.01 | 0.9 | 0.7675 | 7.18 ± 1.94 | 1.2 | 0.3633 | 9.18 ± 1.77 | 1.6 | 0.4251 |
| Chr1.2 | 2.94 ± 1.12 | 2.31 ± 0.78 | 0.8 | 0.1287 | 2.46 ± 1.19 | 0.8 | 0.9299 | 5.02 ± 1.21 | 1.7 | 1.0000 | 7.46 ± 1.98 | 2.5 | 0.9920 | 6.84 ± 0.83 | 2.3 | 0.6902 |
| Chr2 | 3.77 ± 1.63 | 5.72 ± 1.86 | 1.5 | 0.9564 | 2.35 ± 1.58 | 0.6 | 0.9809 | 4.16 ± 1.48 | 1.1 | 0.9983 | 3.64 ± 1.50 | 1.0 | 1.0000 | 11.54 ± 1.40 | 3.1 | 0.0742 |
| Chr3 | 2.29 ± 0.98 | 3.17 ± 1.41 | 1.4 | 0.9980 | 5.80 ± 2.70 | 2.5 | 0.4562 | 7.27 ± 3.49 | 3.2 | 0.4201 | 6.00 ± 1.56 | 2.6 | 0.4289 | 3.80 ± 0.72 | 1.7 | 0.9053 |
| Chr4 | 2.24 ± 0.87 | 7.07 ± 1.16 | 3.2 | 0.3101 | 4.97 ± 0.45 | 2.2 | 0.7340 | 5.85 ± 1.50 | 2.6 | 0.4198 | 13.75 ± 4.23 | 6.1 | 0.0069 | 13.28 ± 3.32 | 5.9 | 0.0090 |
| Chr5 | 4.53 ± 1.93 | 4.45 ± 1.85 | 1.0 | 1.0000 | 13.26 ± 4.01 | 2.9 | 0.0863 | 7.37 ± 2.46 | 1.6 | 0.7349 | 13.73 ± 1.97 | 3.0 | 0.0421 | 13.21 ± 2.11 | 2.9 | 0.0500 |
| Chr6 | 1.60 ± 0.98 | 3.77 ± 1.55 | 2.4 | 0.7973 | 2.85 ± 1.22 | 1.8 | 0.9495 | 4.31 ± 1.68 | 2.7 | 0.5109 | 8.06 ± 3.37 | 5.0 | 0.0654 | 8.29 ± 2.04 | 5.2 | 0.0436 |
| Chr7 | 3.70 ± 1.29 | 3.22 ± 1.24 | 0.9 | 1.0000 | 1.51 ± 0.90 | 0.4 | 0.7279 | 5.09 ± 0.86 | 1.4 | 0.9386 | 7.80 ± 1.21 | 2.1 | 0.3573 | 8.60 ± 2.30 | 2.3 | 0.2542 |
| Chr8 | 5.91 ± 1.29 | 10.13 ± 1.20 | 1.7 | 0.7182 | 12.41 ± 2.23 | 2.1 | 0.3124 | 14.86 ± 4.91 | 2.5 | 0.1643 | 28.10 ± 4.90 | 4.8 | 0.0002 | 34.73 ± 10.20 | 5.9 | <0.0001 |
| Chr9 | 3.92 ± 1.01 | 5.33 ± 1.84 | 1.4 | 0.9921 | 9.76 ± 4.34 | 2.5 | 0.1480 | 10.51 ± 4.60 | 2.7 | 0.2436 | 16.73 ± 3.36 | 4.3 | 0.0041 | 15.30 ± 0.98 | 3.9 | 0.0063 |
| Chr10 | 1.09 ± 0.63 | 4.89 ± 1.66 | 4.5 | 0.2420 | 5.63 ± 1.40 | 5.2 | 0.1632 | 3.93 ± 1.46 | 3.6 | 0.4414 | 5.74 ± 1.33 | 5.3 | 0.1565 | 5.81 ± 1.68 | 5.3 | 0.1482 |
| Chr11 | 8.94 ± 2.77 | 7.46 ± 1.25 | 0.8 | 0.9979 | 16.05 ± 1.77 | 1.8 | 0.2784 | 25.68 ± 5.33 | 2.9 | 0.0052 | 31.80 ± 2.76 | 3.6 | 0.0001 | 35.18 ± 6.74 | 3.9 | <0.0001 |
| Chr12 | 3.22 ± 1.01 | 2.12 ± 1.24 | 0.7 | 0.9872 | 4.78 ± 2.07 | 1.5 | 0.9858 | 5.15 ± 1.51 | 1.6 | 0.8300 | 9.50 ± 2.74 | 3.0 | 0.0796 | 7.48 ± 0.89 | 2.3 | 0.3355 |
| Chr13 | 4.31 ± 1.94 | 5.45 ± 5.45 | 1.3 | 1.0000 | 5.90 ± 1.55 | 1.4 | 0.9614 | 4.61 ± 0.94 | 1.1 | 1.0000 | 4.18 ± 1.72 | 1.0 | 1.0000 | 4.41 ± 1.92 | 1.0 | 1.0000 |
| Chr14 | 4.61 ± 1.34 | 4.55 ± 0.78 | 1.0 | 1.0000 | 14.40 ± 0.89 | 3.1 | 0.0308 | 17.88 ± 2.20 | 3.9 | 0.0047 | 20.73 ± 4.02 | 4.5 | 0.0009 | 37.50 ± 3.58 | 8.1 | <0.0001 |
| Chr15 | 1.57 ± 0.53 | 1.44 ± 0.85 | 0.9 | 1.0000 | 2.88 ± 0.80 | 1.8 | 0.9575 | 2.02 ± 0.76 | 1.3 | 0.9957 | 3.99 ± 1.24 | 2.5 | 0.6720 | 5.12 ± 1.88 | 3.3 | 0.3162 |
| Chr16 | 10.00 ± 1.88 | 7.31 ± 1.70 | 0.7 | 0.9444 | 14.49 ± 3.40 | 1.4 | 0.7512 | 18.50 ± 1.54 | 1.9 | 0.2331 | 20.18 ± 2.70 | 2.0 | 0.1190 | 30.38 ± 6.63 | 3.0 | 0.0020 |
| Chr17 | 6.23 ± 2.19 | 12.11 ± 2.72 | 1.9 | 0.4845 | 15.40 ± 2.14 | 2.5 | 0.0863 | 6.47 ± 1.40 | 1.0 | 1.0000 | 14.82 ± 3.87 | 2.4 | 0.1516 | 34.70 ± 3.78 | 5.6 | <0.0001 |
| Chr18 | 3.87 ± 1.68 | 5.34 ± 0.99 | 1.4 | 0.9899 | 7.78 ± 1.73 | 2.0 | 0.6284 | 6.34 ± 2.01 | 1.6 | 0.8538 | 5.70 ± 0.70 | 1.5 | 0.9361 | 14.53 ± 1.48 | 3.8 | 0.0108 |
| Chr19 | 3.42 ± 0.70 | 3.92 ± 1.65 | 1.1 | 0.9999 | 3.16 ± 1.25 | 0.9 | 1.0000 | 3.64 ± 1.31 | 1.1 | 1.0000 | 9.96 ± 4.14 | 2.9 | 0.1563 | 9.08 ± 2.99 | 2.7 | 0.2403 |

[1] Minimum mutation frequency considering unique mutations within a respective locus.
[2] Fold-change of minimum mutation frequency versus 0 mg/kg/day BbF.
[3] Pairwise comparison to 0 mg/kg/day BbF of the respective locus by GLMM, significant if P < 0.05.

Supplementary Table 6: Minimum frequencies per loci detected in liver samples after exposure to increasing doses of BbF.

| Location of locus | 0 mg/kg/day | 6.25 mg/kg/day | | | 12.5 mg/kg/day | | | 25 mg/kg/day | | | 50 mg/kg/day | | | 100 mg/kg/day | | |
|---|---|---|---|---|---|---|---|---|---|---|---|---|---|---|---|---|
| | $MF_{min}$[1] ± SEM (x $10^{-8}$) | $MF_{min}$[1] ± SEM (x $10^{-8}$) | Fold-change[2] | Adj. P-Value[3] | $MF_{min}$[1] ± SEM (x $10^{-8}$) | Fold-change[2] | Adj. P-Value[3] | $MF_{min}$[1] ± SEM (x $10^{-8}$) | Fold-change[2] | Adj. P-Value[3] | $MF_{min}$[1] ± SEM (x $10^{-8}$) | Fold-change[2] | Adj. P-Value[3] | $MF_{min}$[1] ± SEM (x $10^{-8}$) | Fold-change[2] | Adj. P-Value[3] |
| Chr1 | 2.77 ± 1.20 | 3.96 ± 1.63 | 1.4 | 0.9412 | 8.72 ± 2.22 | 3.1 | 0.0679 | 12.09 ± 2.21 | 4.4 | 0.0147 | 27.15 ± 7.81 | 9.8 | < 0.0001 | 41.48 ± 4.59 | 15.0 | <0.0001 |
| Chr1.2 | 9.80 ± 5.52 | 8.92 ± 1.54 | 0.9 | 1.0000 | 12.93 ± 3.08 | 1.3 | 0.7896 | 23.64 ± 5.74 | 2.4 | 0.0097 | 28.18 ± 5.86 | 2.9 | 0.0010 | 45.28 ± 6.47 | 4.6 | <0.0001 |
| Chr2 | 2.63 ± 1.03 | 2.42 ± 0.84 | 0.9 | 0.9996 | 6.87 ± 1.66 | 2.6 | 0.3570 | 4.78 ± 1.63 | 1.8 | 0.6473 | 17.62 ± 4.60 | 6.7 | 0.0010 | 30.70 ± 8.09 | 11.7 | <0.0001 |
| Chr3 | 3.97 ± 1.79 | 6.18 ± 1.28 | 1.6 | 0.9429 | 7.04 ± 1.25 | 1.8 | 0.9090 | 8.98 ± 1.89 | 2.3 | 0.5024 | 15.88 ± 2.79 | 4.0 | 0.0113 | 35.23 ± 8.91 | 8.9 | <0.0001 |
| Chr4 | 5.59 ± 2.76 | 4.74 ± 2.01 | 0.8 | 0.9993 | 7.01 ± 1.90 | 1.3 | 0.9089 | 15.13 ± 1.96 | 2.7 | 0.0282 | 26.93 ± 2.22 | 4.8 | 0.0001 | 39.43 ± 7.87 | 7.1 | <0.0001 |
| Chr5 | 6.49 ± 1.59 | 5.46 ± 1.86 | 0.8 | 0.9850 | 4.82 ± 1.66 | 0.7 | 0.9964 | 9.39 ± 0.42 | 1.4 | 0.7252 | 17.43 ± 3.19 | 2.7 | 0.0153 | 42.63 ± 5.68 | 6.6 | <0.0001 |
| Chr6 | 5.21 ± 2.90 | 2.00 ± 1.30 | 0.4 | 0.7553 | 4.84 ± 1.06 | 0.9 | 1.0000 | 7.86 ± 0.52 | 1.5 | 0.8318 | 15.32 ± 4.15 | 2.9 | 0.0269 | 32.38 ± 8.14 | 6.2 | <0.0001 |
| Chr7 | 2.26 ± 0.90 | 3.15 ± 1.19 | 1.4 | 0.9871 | 9.21 ± 1.12 | 4.1 | 0.0454 | 11.43 ± 2.15 | 5.1 | 0.0125 | 14.56 ± 4.26 | 6.4 | 0.0016 | 29.85 ± 7.51 | 13.2 | <0.0001 |
| Chr8 | 4.52 ± 1.51 | 4.30 ± 2.59 | 1.0 | 1.0000 | 7.09 ± 1.11 | 1.6 | 0.9638 | 7.72 ± 1.45 | 1.7 | 0.8537 | 27.38 ± 3.01 | 6.1 | 0.0001 | 39.25 ± 6.63 | 8.7 | <0.0001 |
| Chr9 | 6.71 ± 0.73 | 7.92 ± 3.11 | 1.2 | 0.9993 | 12.33 ± 2.56 | 1.8 | 0.2567 | 11.25 ± 2.00 | 1.7 | 0.4410 | 29.98 ± 4.61 | 4.5 | < 0.0001 | 32.43 ± 4.93 | 4.8 | <0.0001 |
| Chr10 | 7.10 ± 1.18 | 5.30 ± 2.78 | 0.7 | 0.9913 | 3.28 ± 0.70 | 0.5 | 0.4860 | 11.61 ± 3.75 | 1.6 | 0.6971 | 11.09 ± 2.05 | 1.6 | 0.7661 | 24.75 ± 3.71 | 3.5 | 0.0014 |
| Chr11 | 10.46 ± 4.19 | 8.65 ± 2.48 | 0.8 | 1.0000 | 6.57 ± 0.99 | 0.6 | 0.8447 | 18.65 ± 1.62 | 1.8 | 0.1140 | 29.75 ± 3.68 | 2.8 | 0.0005 | 37.95 ± 7.50 | 3.6 | <0.0001 |
| Chr12 | 2.12 ± 0.72 | 6.63 ± 1.52 | 3.1 | 0.5879 | 7.94 ± 1.25 | 3.7 | 0.2260 | 9.33 ± 2.56 | 4.4 | 0.0684 | 23.53 ± 5.03 | 11.1 | 0.0001 | 38.20 ± 4.77 | 18.0 | <0.0001 |
| Chr13 | 3.12 ± 1.49 | 3.07 ± 1.84 | 1.0 | 0.9999 | 3.81 ± 1.67 | 1.2 | 0.9844 | 8.84 ± 3.01 | 2.8 | 0.1332 | 9.67 ± 1.18 | 3.1 | 0.0789 | 17.00 ± 1.72 | 5.4 | 0.0013 |
| Chr14 | 10.09 ± 2.96 | 9.40 ± 3.54 | 0.9 | 0.9879 | 9.59 ± 2.01 | 1.0 | 1.0000 | 14.90 ± 3.13 | 1.5 | 0.5698 | 29.23 ± 4.65 | 2.9 | 0.0011 | 63.35 ± 12.50 | 6.3 | <0.0001 |
| Chr15 | 3.16 ± 2.46 | 3.51 ± 2.13 | 1.1 | 0.9964 | 3.77 ± 1.00 | 1.2 | 0.9938 | 7.67 ± 2.81 | 2.4 | 0.3084 | 16.18 ± 1.36 | 5.1 | 0.0019 | 30.10 ± 4.28 | 9.5 | <0.0001 |
| Chr16 | 2.02 ± 0.68 | 6.87 ± 3.22 | 3.4 | 0.4963 | 12.29 ± 2.10 | 6.1 | 0.0225 | 12.86 ± 6.65 | 6.4 | 0.0496 | 24.50 ± 4.35 | 12.1 | 0.0001 | 38.85 ± 3.96 | 19.2 | <0.0001 |
| Chr17 | 4.43 ± 1.85 | 12.65 ± 2.01 | 2.9 | 0.1781 | 12.27 ± 3.76 | 2.8 | 0.1835 | 5.03 ± 1.85 | 1.1 | 0.9972 | 29.05 ± 4.35 | 6.6 | < 0.0001 | 42.50 ± 5.09 | 9.6 | <0.0001 |
| Chr18 | 7.94 ± 3.88 | 8.18 ± 1.92 | 1.0 | 1.0000 | 11.42 ± 2.33 | 1.4 | 0.7965 | 12.58 ± 1.50 | 1.6 | 0.5288 | 33.00 ± 7.80 | 4.2 | < 0.0001 | 44.88 ± 4.12 | 5.7 | <0.0001 |
| Chr19 | 6.78 ± 0.90 | 5.33 ± 1.17 | 0.8 | 0.9893 | 9.62 ± 2.38 | 1.4 | 0.7218 | 8.28 ± 2.36 | 1.2 | 0.9669 | 18.98 ± 3.27 | 2.8 | 0.0123 | 19.98 ± 1.78 | 2.9 | 0.0063 |

[1] Minimum mutation frequency considering unique mutations within a respective locus.
[2] Fold-change of minimum mutation frequency versus 0 mg/kg/day BbF.
[3] Pairwise comparison to 0 mg/kg/day BbF of the respective locus by GLMM, significant if P < 0.05.

Supplementary Table 7: Minimum frequencies of single base substitutions detected in bone marrow and liver samples after exposure to increasing doses of BbF.

| Mutation Subtype | 0 mg/kg/day | 6.25 mg/kg/day | | | 12.5 mg/kg/day | | | 25 mg/kg/day | | | 50 mg/kg/day | | | 100 mg/kg/day | | |
|---|---|---|---|---|---|---|---|---|---|---|---|---|---|---|---|---|
| | $MF_{min}$[1] ± SEM (x $10^{-8}$) | $MF_{min}$[1] ± SEM (x $10^{-8}$) | Fold-change[2] | Adj. P-Value[3] | $MF_{min}$[1] ± SEM (x $10^{-8}$) | Fold-change[2] | Adj. P-Value[3] | $MF_{min}$[1] ± SEM (x $10^{-8}$) | Fold-change[2] | Adj. P-Value[3] | $MF_{min}$[1] ± SEM (x $10^{-8}$) | Fold-change[2] | Adj. P-Value[3] | $MF_{min}$[1] ± SEM (x $10^{-8}$) | Fold-change[2] | Adj. P-Value[3] |
| | | | | | | | | Bone marrow | | | | | | | | |
| C:G>A:T | 2.08 ± 0.40 | 3.28 ± 0.89 | 1.6 | 0.7761 | 5.48 ± 0.76 | 2.6 | 0.0184 | 4.17 ± 0.32 | 2.0 | 0.0912 | 8.84 ± 1.32 | 4.3 | 0.0002 | 12.78 ± 1.41 | 6.1 | <0.0001 |
| C:G>G:C | 0.39 ± 0.23 | 0.86 ± 0.45 | 2.2 | 0.0340 | 1.45 ± 0.16 | 3.7 | 0.0340 | 1.48 ± 0.16 | 3.8 | 0.0523 | 1.61 ± 0.17 | 4.1 | 0.0189 | 2.50 ± 0.29 | 6.4 | 0.0013 |
| C:G>T:A | 2.34 ± 0.24 | 2.79 ± 0.67 | 1.2 | 0.9935 | 2.59 ± 0.64 | 1.1 | 0.9998 | 3.90 ± 0.30 | 1.7 | 0.2643 | 5.63 ± 0.91 | 2.4 | 0.0037 | 7.84 ± 0.81 | 3.4 | 0.0001 |
| T:A>A:T | 0.26 ± 0.13 | 0.45 ± 0.10 | 1.7 | 0.8744 | 1.17 ± 0.16 | 4.5 | 0.0489 | 1.76 ± 0.42 | 6.8 | 0.0103 | 2.04 ± 0.16 | 7.8 | 0.0044 | 2.28 ± 0.56 | 8.8 | 0.0027 |
| T:A>C:G | 0.95 ± 0.15 | 1.44 ± 0.35 | 1.5 | 0.9263 | 1.24 ± 0.33 | 1.3 | 0.8778 | 1.65 ± 0.24 | 1.7 | 0.6433 | 2.15 ± 0.57 | 2.3 | 0.0439 | 1.75 ± 0.43 | 1.8 | 0.2853 |
| T:A>G:C | 0.35 ± 0.05 | 0.61 ± 0.15 | 1.7 | 0.5553 | 0.98 ± 0.11 | 2.8 | 0.0052 | 0.62 ± 0.09 | 1.8 | 0.2107 | 0.81 ± 0.17 | 2.3 | 0.0169 | 0.81 ± 0.11 | 2.3 | 0.0168 |
| Insertion | 0.12 ± 0.08 | 0.10 ± 0.06 | 0.8 | 1.0000 | 0.17 ± 0.07 | 1.4 | 0.9570 | 0.24 ± 0.09 | 2.0 | 0.9409 | 0.32 ± 0.08 | 2.7 | 0.3450 | 0.21 ± 0.07 | 1.8 | 0.7503 |
| Deletion | 0.95 ± 0.17 | 0.70 ± 0.17 | 0.7 | 0.8069 | 0.99 ± 0.05 | 1.0 | 0.9996 | 1.05 ± 0.08 | 1.1 | 0.9949 | 1.28 ± 0.22 | 1.3 | 0.7746 | 1.03 ± 0.26 | 1.1 | 0.9999 |
| MNV[4] | 0.06 ± 0.03 | 0.28 ± 0.14 | 4.6 | 0.5715 | 0.26 ± 0.18 | 4.3 | 0.6452 | 0.19 ± 0.08 | 3.2 | 0.5672 | 0.20 ± 0.05 | 3.3 | 0.6186 | 0.41 ± 0.04 | 6.8 | 0.1553 |
| | | | | | | | | Liver | | | | | | | | |
| C:G>A:T | 1.21 ± 0.29 | 2.53 ± 0.46 | 2.1 | 0.8117 | 3.85 ± 0.15 | 3.2 | 0.4030 | 9.03 ± 0.76 | 7.5 | 0.0319 | 21.38 ± 4.07 | 17.7 | 0.0009 | 35.48 ± 5.62 | 29.3 | 0.0001 |
| C:G>G:C | 0.38 ± 0.20 | 0.44 ± 0.20 | 1.2 | 1.0000 | 1.26 ± 0.33 | 3.3 | 0.4026 | 2.31 ± 0.69 | 6.1 | 0.0541 | 4.56 ± 0.79 | 12.0 | 0.0055 | 7.77 ± 0.94 | 20.4 | 0.0007 |
| C:G>T:A | 4.75 ± 0.57 | 4.88 ± 1.29 | 1.0 | 0.9910 | 5.53 ± 0.75 | 1.2 | 0.9015 | 5.34 ± 0.57 | 1.1 | 0.9964 | 10.41 ± 0.85 | 2.2 | 0.0043 | 16.53 ± 1.37 | 3.5 | <0.0001 |
| T:A>A:T | 0.87 ± 0.18 | 0.75 ± 0.33 | 0.9 | 0.9617 | 1.36 ± 0.34 | 1.6 | 0.8657 | 2.47 ± 0.21 | 2.8 | 0.1372 | 4.42 ± 0.55 | 5.1 | 0.0034 | 7.52 ± 1.04 | 8.6 | 0.0034 |
| T:A>C:G | 1.05 ± 0.30 | 1.00 ± 0.38 | 1.0 | 0.9935 | 1.54 ± 0.11 | 1.5 | 0.6879 | 1.32 ± 0.09 | 1.3 | 0.9461 | 2.22 ± 0.33 | 2.1 | 0.0909 | 3.47 ± 0.51 | 3.3 | 0.0016 |
| T:A>G:C | 0.40 ± 0.17 | 0.65 ± 0.14 | 1.6 | 0.8476 | 0.62 ± 0.08 | 1.6 | 0.8725 | 0.80 ± 0.30 | 2.0 | 0.6951 | 0.96 ± 0.1 | 2.4 | 0.3025 | 1.66 ± 0.32 | 4.2 | 0.0137 |
| Insertion | 0.24 ± 0.07 | 0.12 ± 0.05 | 0.5 | 0.9145 | 0.36 ± 0.16 | 1.5 | 0.8505 | 0.23 ± 0.04 | 1.0 | 1.0000 | 0.25 ± 0.1 | 1.0 | 0.9986 | 0.45 ± 0.03 | 1.9 | 0.5266 |
| Deletion | 0.81 ± 0.27 | 0.67 ± 0.16 | 0.8 | 1.0000 | 0.76 ± 0.07 | 0.9 | 1.0000 | 0.67 ± 0.23 | 0.8 | 0.9999 | 1.00 ± 0.13 | 1.2 | 0.9244 | 1.65 ± 0.33 | 2.0 | 0.0609 |
| MNV[4] | 0.14 ± 0.14 | 0.15 ± 0.01 | 1.1 | 0.9975 | 0.20 ± 0.04 | 1.4 | 0.9325 | 0.39 ± 0.08 | 2.8 | 0.2172 | 0.68 ± 0.09 | 4.9 | 0.0298 | 1.06 ± 0.09 | 7.6 | 0.0039 |

[1] Minimum mutation frequency considering unique mutations.
[2] Fold-change of minimum mutation frequency versus 0 mg/kg/day BbF.
[3] Pairwise comparison to 0 mg/kg/day BbF by GLMM, significant if P < 0.05.
[4] Multi-nucleotide variants.

Supplementary Table 8: Benchmark doses of measured genotoxic endpoints in blood, bone marrow and liver.

| Genotoxic endpoint | n | CES[1] | BMDL[2] | BMDU[3] | Log10 BMDL | Log10 BMDU | Bootstrap runs |
|---|---|---|---|---|---|---|---|
| MN RET | 8 | 0.5 | 0.408 | 4.06 | -0.3893 | 0.6085 | 200 |
| MN RBC | 8 | 0.5 | 2.23 | 9.34 | 0.3483 | 0.9703 | 200 |
| MF BM | 4 | 0.5 | 3.6 | 12.8 | 0.5563 | 1.1072 | 200 |
| MF Liver | 4 | 0.5 | 10.1 | 18.4 | 1.0043 | 1.2648 | 200 |

[1]Critical effect size used to perform benchmark dose modeling.
[2]Benchmark dose lower confidence limit.
[3]Benchmark dose upper confidence limit.

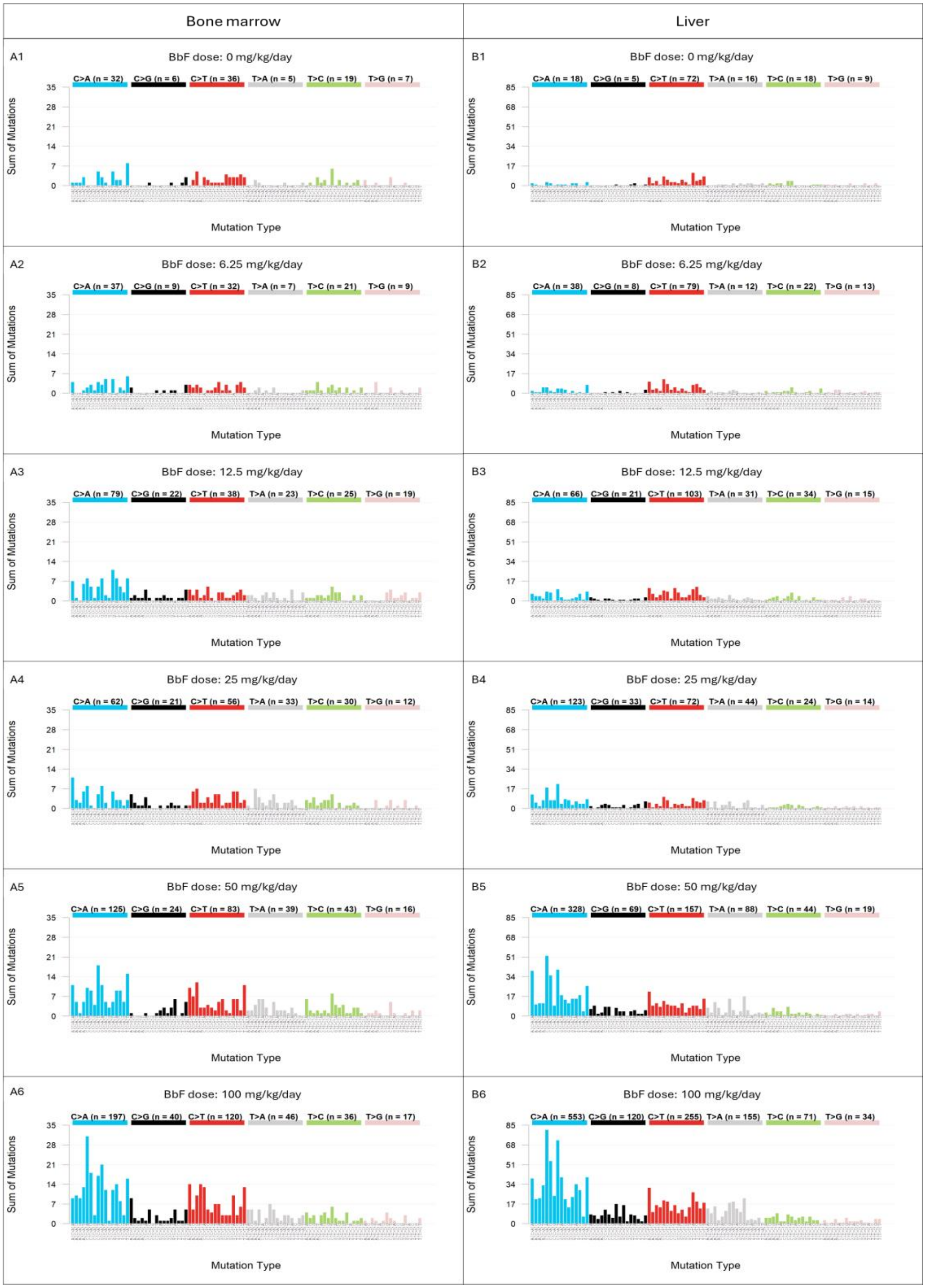


Supplementary Figure S1: Trinucleotide mutation pattern of bone marrow (A) and liver (B) samples after 28-day exposure to the vehicle control (A1, B1) and increasing doses of BbF (A2 = 6.25, A3 = 12.5, A4 = 25, A5 = 50, A6 = 100, B2 = 6.25, B3 = 12.5, B4 = 25, B5 = 50 and B6 = 100 mg/kg/day). The panels represent the sum of mutations of each 96-mutational subtypes for n = 4 samples of each dose group. The sum of all mutation for each single base substitution type; C>A (blue), C>G (black), C>T (red), T>A (gray), T>C (green) and T>G (pink), are indicated.

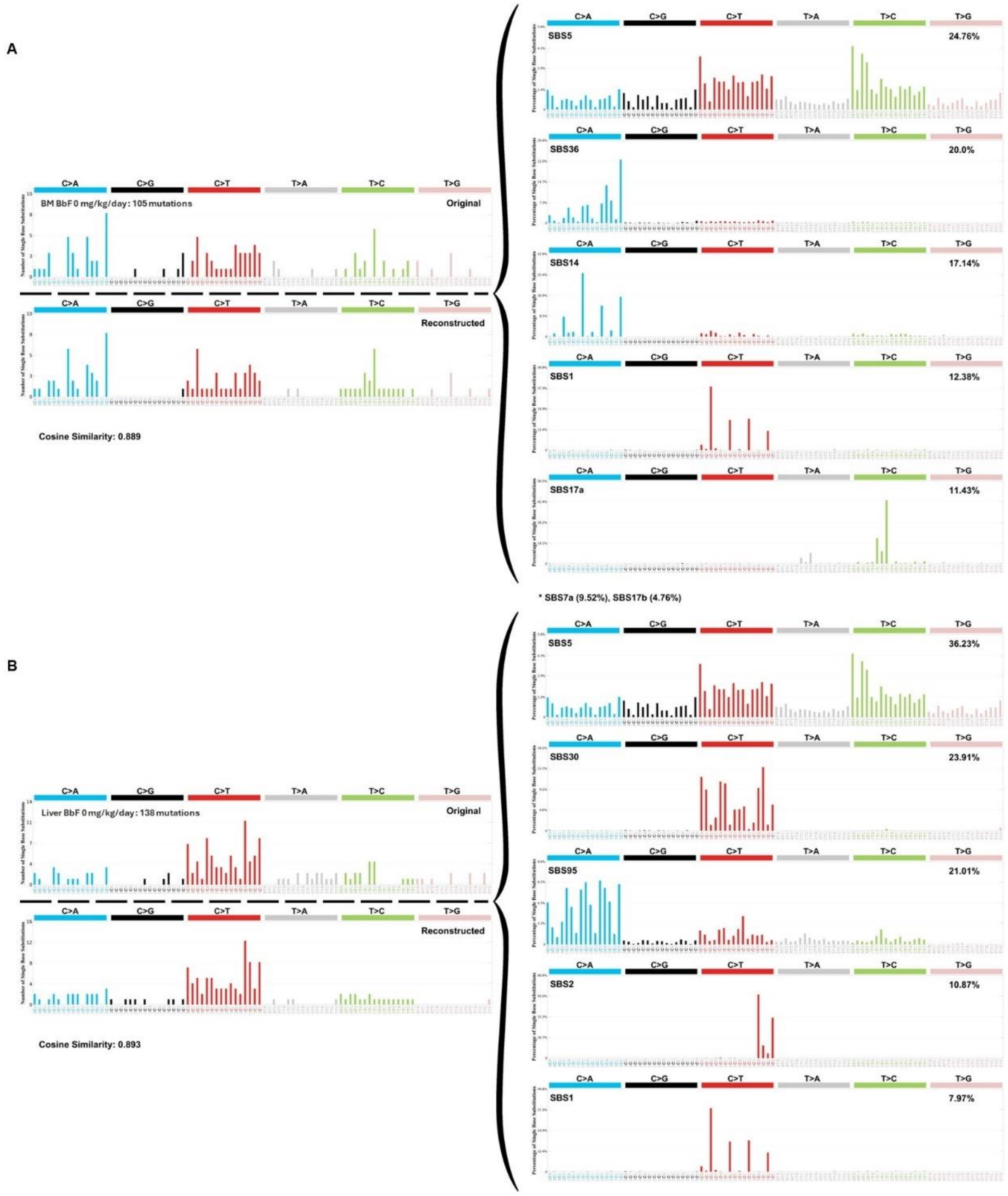


Supplementary Figure 2. Mutation signature analyses for vehicle control in bone marrow (A) and liver (B). The original trinucleotide mutation profile is shown on the top left. SigProfilerAssignment used the single base substitution (SBS) signatures of the Catalogue of Somatic Mutations in Cancer (COSMIC) database to reconstruct the signatures. The SBS signatures and their relative contributions are shown on the right. The reconstructed mutation pattern, number of substitutions used for the assignment and cosine similarity between reconstructed and observed trinucleotide mutation profile are shown on left below the original profile. The total amount of mutations of the original signature are indicated on the left.
